# The effect of the consumption of toxic prey on avian predator physiology

**DOI:** 10.64898/2026.10.14.759177

**Authors:** Oihana Garcia Carrillo, Elsa Minoza, Christopher J. Irving, Suvi Ruuskanen, Hannu Pakkanen, Liisa Hämäläinen

## Abstract

Many insects sequester plant toxic metabolites as a chemical defence against predators. Predators are predicted to include chemically defended prey in their diet if the benefits of gaining nutrients outweigh the costs of ingesting toxins. However, the physiological costs associated with the consumption of prey toxins remain largely unknown, especially when using ecologically relevant predators and naturally occurring levels of toxins. Here, we investigated whether the consumption of chemically-defended wood tiger moths (*Arctia plantaginis*) influences the physiological state of an avian predator, the great tit (*Parus major*). We manipulated the moth toxicity by first rearing the larvae in two different diets that varied in pyrrolizidine alkaloid (PA) content, and then manually extracting part of the defensive secretions from half of the moths in both diet groups. This led to four treatments with different moth PA profiles, which were fed to great tits for six consecutive days. After the bird diet manipulation, we collected fecal and blood samples for various physiological analyses of stress and metabolism. We found no significant effects of PA consumption on corticosterone or serotonin concentrations, nor on oxidative stress biomarkers. However, birds exposed to higher PA doses used more of their mitochondrial maximal capacity, which might result from increased metabolic stress. Overall, our results suggest that incorporating some PA-defended insects in the diet does not incur high physiological costs to avian predators, but further studies are needed to explore the costs of consuming a higher number of defended prey and the interactive effects of different prey toxins.

## INTRODUCTION

Predation is a major driver of evolution, shaping selection for various mechanisms to counter and minimise predator attacks (Ruxton et al. 2018). Aposematism is a widespread antipredatory defence, where prey advertise their unprofitability through conspicuous warning signals (Poulton 1890; Ruxton et al. 2018). A prime example of warningly colored prey is chemically defended insects that often sequester and utilise plants’ toxic secondary metabolites as a defence against predators and parasitoids (Opitz and Müller 2009). Despite these chemical defences, predators can attack and consume aposematic prey, especially if the alternative prey is scarce or the predators are in a poor energetic state (Sherratt 2003; Barnett et al. 2007, 2012). These decisions to attack chemically defended prey are predicted to depend on the nutritional value of the prey and the physiological costs of ingesting prey toxins (Skelhorn et al. 2016). Accordingly, prior toxin exposure (toxin load) has been shown to reduce predator attacks on chemically defended prey (Skelhorn and Rowe 2007), suggesting that predators actively regulate their toxin intake. However, the specific physiological effects of toxin consumption that mediate a predator’s future foraging decisions remain poorly understood.

One common group of chemical defences found in many insects are pyrrolizidine alkaloids (PAs) that many aposematic species, such as Arctiinae moths, sequester from plants (Rothschild et al. 1979; Von Nickisch-Rosenegk and Wink 1993; Trigo 2011). PAs in plants and insects commonly co-exist with their corresponding N-oxides which are less toxic but can be metabolically reduced back into their free base form when ingested (Nuringtyas et al. 2014; Yang et al. 2019). Consuming PA-defended prey elicits aversion responses from predators (Rowell-Rahier et al. 1995; Cardoso 1997), but the research on the mechanistic effects of PAs has mostly focused on humans and livestock consuming PA-containing plants (reviewed in Wiedenfeld and Edgar 2011). The metabolic pathway underlying the toxic effects includes dehydrogenation of PAs in the liver to generate dehydropyrrolizidine alkaloids (DHPAs), which may further bind to cellular proteins to form pyrrole-protein adducts, leading to hepatotoxicity (Xu et al. 2019; He et al. 2021). Interestingly, PA susceptibility seems to vary highly among species, which is suggested to result from differences in both the metabolic activation to form DHPAs, as well as the detoxification pathways to convert PAs to less reactive metabolites (He et al. 2021). Birds are an important predator group for many aposematic insects, but previous research with birds has mainly investigated the toxicity of plant PAs on poultry (Buckmaster et al. 1977; Cheeke and Pierson-Goeger 1983; Diaz et al. 2014; Brown et al. 2016). These studies have shown that high doses of ingested PAs can have detrimental effects on production traits, such as growth and egg production in laying hens (*Gallus gallus domesticus*, Eröksüz et al. 2003; Diaz et al. 2014) and japanese quails (*Coturnix japonica*, Eröksüz et al. 2008), and often lead to liver damage (Brown et al. 2016). However, other potential physiological costs remain poorly understood and are likely to vary among species. Examining detoxification costs and physiological effects of naturally occurring prey PA levels on ecologically relevant wild avian predators is therefore necessary to understand predators’ adaptive foraging decisions and the resulting selection pressures for prey chemical defences.

One potential physiological cost of consuming chemically defended prey is increased stress and changes in animal metabolism (Harvey et al. 1994). During a stressful event, glucocorticoids are rapidly secreted and can be sustained for prolonged periods (Sapolsky et al. 2000). In birds, corticosterone is the primary glucocorticoid, which is produced in response to the stimulation of the adrenal cortex by adrenocorticotropic hormones (Hess 2002). Its release is a coping mechanism to facilitate a shift in energy balance (Sapolsky et al. 2000), which could be potentially related to the consumption of prey toxins, as PAs can cause hepatotoxicity (Xu et al. 2019) that could lead to metabolic stress and the activation of the hypothalamic-pituitary-adrenal (HPA) axis. This makes the measure of corticosterone levels relevant as a proxy of stress that may be induced by the consumption of chemically defended prey.

At the cellular level, PA consumption has been shown to induce oxidative stress in mammals (Xu et al. 2019). Oxidative stress refers to an imbalance between the production of reactive oxygen species (ROS) and the organism’s antioxidant defences, leading to an accumulation of cellular damage to lipids, proteins and DNA (Halliwell and Gutteridge 2015). PAs have been demonstrated to disrupt the normal cellular redox status when toxic reactive metabolites react with glutathione (GSH), an important antioxidant that contributes to PA detoxification (reviewed in Xu et al. 2019). This may lead to depletion of GSH and overproduction of ROS and is often linked to changes in the activity of antioxidant enzymes such as glutathione-S-transferases (GSTs), which are involved in the detoxification of ROS (Fu et al. 2004; Liang et al. 2009). The resulting imbalance of the antioxidant system increases oxidative stress, which may further lead to PA-induced hepatotoxicity (Xu et al. 2019). The same PAs used in these toxicity studies are commonly found in chemically defended insects (e.g. senecionine, Trigo 2011), but whether the concentrations found in insects are high enough to induce oxidative stress for avian predators remains unknown. If so, this could have important fitness effects, as oxidative stress can affect both longevity and reproduction in birds (Costantini 2008).

Increased corticosterone levels or oxidative stress could further affect energetics at the cellular level by directly impairing mitochondrial respiration (Cardoso et al. 1999; Manoli et al. 2007). In aerobic organisms, mitochondria are the main source of cellular adenosine triphosphate (ATP) production through oxidative phosphorylation (OXPHOS), which can play an important role in an individual’s fitness (Spinelli and Haigis 2018). Mitochondria are highly dynamic organelles that respond to changing cellular physiology, nutrient availability and energy demands (Spinelli and Haigis 2018). Thus, changes in diet composition, such as the ingestion of PAs, might impact mitochondrial respiration. Previous in vivo studies using high PA concentrations on rat liver cells suggest that PAs can impair mitochondrial function (Mingatto et al. 2007; Lu et al. 2018); however, whether such effect occurs beyond mammals and with ecologically relevant toxin levels is unclear.

The consumption of prey toxins could also affect individuals via stress-independent molecular mechanisms by affecting predators’ serotonin levels. Serotonin (5-hydroxytryptamine, 5-HT) is a monoamine neurotransmitter, implicated in a wide range of physiological processes in vertebrates (Mohammad-Zadeh et al. 2008). Previous research suggests that PA molecules could have an agonistic action on serotonin receptors (Schmeller et al. 1997), but again, it is unknown if similar effects are seen in predators that consume insects defended with PAs in lower concentrations. Investigating whether the consumption of PAs influences serotonin levels of avian predators is particularly interesting because serotonergic system is linked to many behaviors, such as aggression, risk-taking behavior and feeding (Timm et al. 2018; Saldanha et al. 2023), and changes in serotonin levels could therefore further impact predator foraging decisions and predator-prey interactions.

In this experiment, we aimed to explore the physiological effects of ecologically relevant levels of prey toxins on an avian predator, the great tit (*Parus major*). Great tits are an ideal model species to answer these questions, as they are commonly used in physiology (e.g. Rainio et al. 2015) and predator-prey research (e.g. Gordon et al. 2021). They are generalist predators, and during the summer their diet mainly consists of different insect species (Royama 1970), so they are potential predators for many chemically defended insects. The model prey in the experiment was the aposematic wood tiger moth (*Arctia plantaginis*), which sequesters PAs from the diet (Winters et al. 2021). Great tits are known to consume wood tiger moths in captive studies (Gordon et al. 2021), and they are also suggested to be one of the main predators of the moths in the wild (Nokelainen et al. 2014).

We first manipulated the toxicity of *A. plantaginis* by rearing larvae in two different diets that varied in PA content, and by removing part of the defensive secretions from half of the adult moths, leading to four groups with different moth PA profiles. These were fed to wild-caught great tits for six consecutive days. After this diet manipulation, we measured the effect of the consumption of moths with different PA levels on several physiological parameters. These included corticosterone levels measured from fecal samples and several oxidative stress markers, mitochondrial metabolism and serotonin levels measured from blood samples. We hypothesised that birds that consumed moths with higher concentrations of PAs would have higher levels of corticosterone and, as observed in other species, higher levels of oxidative stress, as higher toxicity is assumed to translate as a more stressful experience (Xu et al. 2019). As reported in previous experiments (Mingatto et al. 2007; Lu et al. 2018), we also expected mitochondrial metabolism to be affected by toxins, with birds with a higher toxin load having a decreased mitochondrial respiration rate. Finally, we expected birds fed with a higher PA concentration to have lower serotonin concentrations, as PA molecules have been previously reported to interact with serotonin receptors (Schmeller et al. 1997). Since the experiment included treatments with different PA concentrations, the effects were expected to be strongest in the treatment with the highest concentrations of PAs. Elucidating these questions allows us to understand the costs and consequences of ingesting toxic prey for a predator, and thus, are relevant to comprehend the evolution of complex and costly anti-predatory mechanisms such as aposematism.

## METHODS

### Prey species and toxicity manipulation

The aposematic wood tiger moth *A. plantaginis* (Lepidoptera: Erebidae: Arctiinae) is widely distributed in the temperate region of the northern hemisphere. The moths sequester PAs from chemically defended plants (such as groundsel, *Senecio vulgaris*), which makes them unpalatable for predators (Von Nickisch-Rosenegk and Wink 1993; Winters et al. 2021). In addition to chemically sequestered PAs, the species synthesises *de novo* methoxypyrazines, which are released from their prothoracic region in response to danger (Burdfield-Steel et al. 2018). These “neck fluids” (NF) also contain some of the acquired PAs (A. Winters et al. unpublished).

In this experiment, the toxicity of the moths was manipulated by both restricting the larval diet to non-PA-containing plants and by manually removing the defensive neck fluids from adult moths. Twenty pairs of *A. plantaginis* were collected during the summer of 2024 in multiple sites in Central Finland. These parent *A.plantaginis* produced around 2,200 larvae, of which 1,200 were divided into family groups of 10 larvae and housed in plastic containers (12.5 × 9.5 × 10.5 cm). This F1 generation was reared in a greenhouse on two different diets: 1) a PA diet, containing a mixture of groundsel (containing PAs, Cheng et al. 2017) and dandelion (*Taraxacum sp*., not known to contain PAs), and 2) an exclusive dandelion diet (control). The larvae were provided with fresh leaves every day. The greenhouse was set at a constant 21 °C and had UV Lights (Osram Powerstar 400W/D Pro Daylight E40) set from 0500 to 2300 every day, simulating summertime in Central Finland. Once the larvae pupated, they were moved to individual cylindrical containers (height: 6.5cm, diameter: 3.5 cm). When the adults emerged from the pupae (PA: 434 adults, control: 417 adults), their sex, weight and, if male, color morph (white or yellow) was recorded. We found an effect of diet on larval development, with the larvae reared on the PA diet developing more quickly and having higher pupal weight than the larvae reared on the control diet (see appendix).

All moths were given water once they emerged and after this an hour to drink and generate the neck fluids. Then, half of the moths from each treatment group had their neck fluid removed by gently squeezing the thorax behind the head, which stimulated reflex bleeding (Burdfield-Steel et al. 2018). This was repeated until the moth did not release additional fluid. While this technique can not assure that the neck fluid is completely removed from the individual, the toxin levels will be comparatively different to the moths that had their neck fluid retained. This resulted in four treatments: 1) control diet and neck fluid removed (C + no NF), 2) control diet and neck fluid intact (C + NF), 3) PA diet and neck fluid removed (PA + no NF), and 4) PA diet and neck fluid intact (PA + NF). The intention was to create a gradient of toxicity, where C+ no NF would have the lowest toxicity (acting as a control), and PA + NF the highest.

Adults were freeze-killed on the day of emerging and later freeze-dried using a Christ Alpha 1-4 LD Plus Lyophilizer. Individuals from the same treatment (around 170 per treatment) were then blended together and broken down into powder that could be given to birds in standardized doses. Each treatment group had the same proportion of females and males (1:1). The male moths have two color morphs (yellow and white) that might differ in toxicity (Winters et al. 2021), and while the white morph was more common in our lab population, we made sure that the proportion of the two morphs was same in each treatment (15 yellow and 70 white morphs). The powders were stored in a −80°C freezer in air-tight containers to prevent degradation during the experiment.

To quantify the PA concentrations in each treatment group, we extracted PAs from the homogenized moth powders following the protocol described in Reinwaldt et al. (2017). We then used LC-MS/MS to quantify the concentration of the five previously identified PAs in wood tiger moths (retrorsine, seneciphylline, seneciphylline N-oxide, senecionine, senecionine N-oxide, Winters et al. 2021) in each of the four treatments (see appendix for a more detailed description of the PA extraction and the LC-MS/MS protocols). The extractions (three technical replicates from each treatment group) and quantifications were performed at the beginning (t1) and at the end of the four-month bird experiment (t4) to investigate possible degradation in PA concentrations in the frozen moth powder (i.e. storage time effects) and ensure the birds were getting standardized concentrations of toxin. Additionally, at t4 we re-measured the PA concentrations of the extracts from t1. For the C + NF samples, we excluded the results from the quantification at t1, as they were inconsistent with the other two measures and likely to be a measurement error. Otherwise, the three technical replicates within the same quantification time (mean CV% = 17.16), the two quantifications of the same extracts (mean CV% = 15.39) and the two extractions (i.e., storage time effects, mean CV% = 23.45) gave similar results. Although there was some variation in the exact values, the relative concentrations among the treatments remained the same, with PA + NF treatment having the overall highest PA concentrations, and C + no NF the lowest PA concentrations (Table 1, appendix Fig. A2).

**Table 1.** Concentrations of different PAs for each treatment (μg/mg). The values are averages of the three quantifications that each had three technical replicates, except for C + NF where the first quantification was excluded (see the text).

|  | Retrorsine | Seneciphylline | Seneciphylline<br>N-oxide | Senecionine | Senecionine<br>N-oxide |
| --- | --- | --- | --- | --- | --- |
| C + no NF | 0.095×10<sup>-2</sup> | 0.007 | 0.014 | 0.014 | 0.026 |
| C + NF | 0.061×10<sup>-2</sup> | 0.002 | 0.040 | 0.005 | 0.074 |
| PA + no NF | 0.002 | 0.021 | 1.669 | 0.037 | 3.003 |
| PA + NF | 0.009 | 0.176 | 1.554 | 0.263 | 3.101 |

### Predators and housing

We used wild-caught great tits as a model predator to study the physiological effects of the consumption of the chemically defended moths. Experiments took place at Konnevesi research station in Central Finland from November 2024 to March 2025. A total of 80 wild great tits (23 juvenile males, 28 juvenile females, 21 adult males, 8 adult females) were caught from feeding sites. Each bird was sexed and aged (juveniles: < 1 year old, adults: > 1 year old) based on plumage (Svensson 1992). They were weighed and measured (tarsus, wing length) after capture and before releasing, and their body condition index was calculated using capture weight (0.01g) and tarsus length (0.01 cm) measures (Peig and Green 2009). This was assumed to be related to the individual’s health by indicating their energetic reserves. The birds were housed individually in plywood cages (120 × 90 × 60 cm) with enrichment, for approximately 20 days. They were kept at room temperature with a daily light cycle of nine hours. Sunflower seeds were provided *ad libitum*, except before and during diet manipulation, to ensure birds’ motivation to forage. Fresh water was always available. After the experiment, the birds were ringed and released back at the capture site, except for two birds that unfortunately died in captivity for unknown reasons. Wild birds were used with the permits from the Finnish National Project Authorisation Board (ESAVI/34419/2024) and the Centre for Economic Development, Transport and Environment (VARELY/4950/2024).

### Predator diet manipulation and sample collection

In the diet manipulation, birds were divided into four treatments (n = 20 each) that received moths of varying toxicity (see above): 1) C + no NF, 2) C + NF, 3) PA + no NF, and 4) PA + NF. The diet manipulation began on the second day of captivity, and lasted for six days (Fig. 1). On each day, the birds were fed with 54mg of the freeze-dried moth powder, which corresponded to the average weight of one moth, calculated by averaging the dry weight of all moths reared in the colony. Therefore, during the six-day manipulation, birds consumed approximately six moths, which was assumed to be similar to what they might consume in the wild. To encourage birds to eat the moth powder, it was mixed with 470 mg of a dough-like mixture containing dried mealworm and oat powder (112 mg each), peanut butter (22 mg) and approximately 0.2 mL of water to create “a moth pellet”. On the day of the capture, birds were given this mixture without the moth powder to familiarise them with the pellet. During the manipulation, regular food was removed to encourage the birds to eat the moth pellet, and the birds were checked approximately every 30 minutes. The time needed for the birds to eat the entire pellet was recorded (except for the first 16 birds). If the bird did not eat any of the pellet in two hours, it was given 2–3 sunflower seeds to ensure that the food restriction was not too long. Six birds (one in the C + NF treatment, three in the PA no NF treatment and two in the PA + NF treatment) did not consume the entire pellet during the first day, but consumed the pellets on the following days. If the bird refused to eat the pellet also on the second day, it was excluded from the experiment (seven birds).

**Figure 1.**
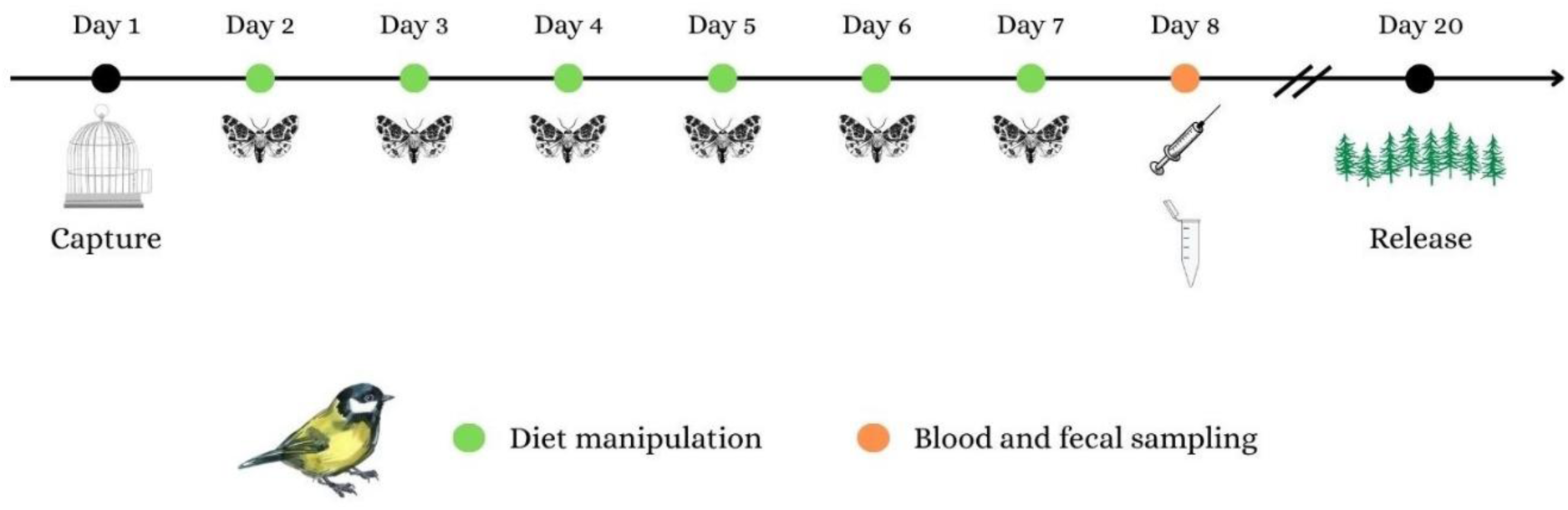
Timeline of the experiment. Day 1: capture. Day 2–7: diet manipulation with moth pellets. Day 8: blood and fecal sampling. Day 20: release. The birds were kept in captivity 12 days after the sampling to conduct behavioral assays related to a simultaneous experiment.

Blood and fecal samples were collected a day after the six-day diet manipulation (see Fig. 1). Fresh fecal samples (approximately 1500 mg) were collected from the bottom of the birds’ cage with tweezers and stored at −80°C until corticosterone extraction. Blood samples (a maximum volume of 120 µL) were taken from the brachial vein, using a needle (MicrolanceTM), and then collected using heparinised capillary tubes (Hirschmann™). From this sample, 30 µL was immediately immersed in liquid nitrogen and stored in a freezer at −80°C for oxidative stress assays, and 20-30 μL was stored at +5°C until mitochondrial respiration measurements. In addition, 10-15 μL of plasma was extracted from the whole blood by centrifuging the sample and stored at −20°C for serotonin analysis. In some cases, we did not get a sufficient amount of blood for each analysis, so the final sample sizes for each physiological assay varied (see below). We also weighed the birds after the diet manipulation to investigate potential effects of PAs on body weight; however, this measure was recorded only from half of the birds (n = 40).

### Physiological measurements

#### Corticosterone concentration

Frozen fecal samples from each treatment (n = 67) were extracted following instructions from Arbor Assays, previously validated for birds (Leach et al. 2024). Samples were analyzed using the Arbor Assays DetectX Corticosterone EIA kit (Ann Arbor, MI, USA). All samples were analyzed in duplicates (mean CV% = 5.10 after excluding 8 birds with CV% > 15) using the Tecan Spark multimode microplate reader at 450nm.

#### Oxidative stress

Oxidative stress was analyzed from blood samples using several commonly used biomarkers: i) the reduced glutathione (GSH) and the reduced/oxidized glutathione ratio (GSH:GSSG), ii) glutathione S-transferases (GST) activity, and iii) lipid peroxidation (using malondialdehyde, MDA, as a proxy), measured via thiobarbituric acid reactive substances (TBARS assay). The GSH:GSSG ratio is an indicator of cellular redox status, reflecting the organism’s ability to neutralise oxidative stress (Isaksson et al. 2005; Rainio et al. 2015). GSH is an antioxidant, while GSSG is the product of its reaction with ROS, so a low GSH:GSSG ratio suggests excessive oxidation and oxidative stress, while a high ratio indicates a good antioxidant defence capacity. GSTs are enzymes involved in the detoxification of ROS and have a central role in the antioxidant defence of cells (Halliwell and Gutteridge 2015). The formation of TBARS is an indirect indicator of cellular damage, as it reflects the degree of lipid degradation by ROS (Espin et al. 2017).

We first prepared samples for each assay by diluting 15 µL of each blood sample with 200 µL of 0.9% NaCl. Protein concentration was measured using bovine serum albumin (BSA) as the standard in a 96-well plate using Bio-Rad Protein Assay Dye (Bio-Rad Laboratories, Cat #500–0006). Once the protein concentrations had been measured and validated, each sample was divided into three tubes for glutathione, GST and TBARS, which were then immersed in liquid nitrogen before being stored at −80°C for future use.

The GSH:GSSG ratio was estimated using the Arbor Assays DetectX Glutathione Fluorescent Detection Kit (Ann Arbor, MI, USA). Before starting, the samples were prepared following kit instructions: 12 µL of blood from each bird (n = 66) was mixed with 48 µL of 5% SSA (Sulfosalicylic Acid + MQ-H2O). The mixture was incubated for 10 minutes on ice allowing the proteins to precipitate and centrifuged for 10 min at 10000 g at 4°C. 30 µL of supernatant was collected and transferred to new tubes and further diluted 1:5 in Assay Buffer (100 mM Na-phosphate buffer and 5 mM EDTA, pH 7.5) to give a final SSA concentration of 1%. Samples were then analyzed in duplicates following the manufacturer’s instructions, estimating the amount of free (GSH, nmol/mg of protein: mean CV% = 3.14, after excluding 3 birds with CV% > 15) and oxidized glutathione (GSSG: mean CV% = 3.12 after excluding 5 birds with CV% > 15). One outlier was removed from the GSH:GSSG ratio analysis based on the Grubb’s test (*p* < 0.001).

GST activity was analyzed following in-house methods as in Rainio et al. (2015). In short, 2 µL of the prepared blood sample (n = 65) was pipetted in triplicate into the wells of a 384-well plate, and 48 µL of reaction mix (Dulbecco’s phosphate buffered Saline, 100 mM 1-Chloro-2,4-dinitrobenzene and 200mM GSH) was added to each well. The absorbance was read at 340nm for 25 times (approximately every 14.7 seconds) and the GST activity was expressed as pmol/min/mg of protein. In most cases (n = 55), we analyzed samples in duplicates (instead of triplicates) to improve CV%. Samples with CV% > 15 in duplicate measurements (n = 19) were excluded. We also excluded samples where the coefficient of determination (R²) in the absorbance changes was < 0.9 in both sample replicates (n = 4). Samples where one of the replicates had slightly lower R² (> 0.83, n = 5) were included. One outlier with high GST value was removed from the analysis based on the Grubb’s test (*p* < 0.001), resulting in a final sample size of 41 birds (mean CV% = 6.79). TBARS analysis (n = 62) was conducted following Espin et al. (2017) using fluorescence and malondealdehyde standard. Each sample was plated in triplicate (mean CV% = 5.64 after excluding 3 birds with CV% > 15), and the MDA concentrations were calculated as nmol/mg of protein.

#### Mitochondrial metabolism

To test the impact of PA toxicity on mitochondrial respiration, we measured mitochondrial aerobic metabolism in each treatment group from January to March (n = 43). The measurements were done within 60 hours from blood sampling to avoid cell deterioration and ensure reliable estimates of the respiration rate. Mitochondrial respiration was analyzed using high-resolution respirometry (Oroboros Instruments, Innsbruck, Austria) at 41°C, adapted from a protocol described in Stier et al. (2019). We used 20 μl to 30 μl of fresh blood, depending on availability, and suspended it in the Mir05 buffer. Three distinct respiration rates were measured: 1) endogenous cellular respiration rate before permeabilisation (ROUTINE), 2) maximum respiration rate fuelled with exogenous substrates of complexes I and II, as well as ADP (CI+II), and 3) the respiration rate supporting ATP synthesis through oxidative phosphorylation (OXPHOS). CI+II alone was not analyzed, but it was used to calculate a mitochondrial flux ratio (FCR) that reflects the proportion of maximal respiration capacity being used under the endogenous cellular condition: FCR ROUTINE/(CI+II). This indicates the cellular control of the mitochondrial respiration by endogenous ATP turnover and substrate availability. The values were extracted from the raw data twice (by the same observer) with individuals in random order to avoid bias and ensure repeatability (mean CV% = 1.7). The calculations were standardized based on the volume of blood used. In addition, the number of cells in each sample was measured using the TC20 Automated Cell Counter | Bio-Rad cell counter, and this value was taken into account in the statistical analyses.

#### Serotonin concentration

Serotonin (n = 42) was analyzed using a commercially available Serotonin ELISA kit (ADI-900-175, Enzo Ltd., Farmingdale, NY), following the manufacturer’s instructions. Most samples were analyzed in duplicates (n = 29, mean CV% = 6.55 after excluding four birds with CV% > 18), but we also used single measurements if there was not enough plasma for duplicates (n = 13). More detailed descriptions of the protocols for the physiological assays are included in appendix.

### Statistical analyses

We tested the effect of the moth’s toxicity on birds’ behavior and physiological measures using linear and linear mixed effects models with a Gaussian error distribution. All models included bird diet treatment as a four-level categorical variable (C + no NF/ C + NF/ PA + no NF/ PA + NF). We started model selections with models that also included birds’ sex, age (adult/juvenile) and body condition, as well as other variables relevant for each analysis (see below). Terms in the final models were selected based on the Akaike Information Criterion (AIC) and statistical significance. Other variables than treatment were removed if they were not significant (threshold α = 0.05) and did not improve the AIC (see model selections in appendix). The final sample size for each analysis varied depending on the availability of samples for each assay (Table A2 in appendix). To increase statistical power to detect differences between birds that received low and high PA concentrations (Table 1), we also ran separate models for each response variable including a moth diet (PA/ no PA) as a fixed effect instead of the four separate treatment levels. This was done by combining PA + NF and PA + no NF treatments into one “PA treatment”, and C + NF and C no NF treatments into one “control treatment”, which increased the sample size per treatment.

To analyze whether bird diet treatment influenced the time required to consume the moth pellets, we used a linear mixed effects model with the time to eat the pellets (in minutes) as a response variable (n = 65). In addition to the bird diet treatment, fixed effects in the final model included the day of bird diet manipulation (1–6 as a continuous variable), and bird identity was included as a random effect to control for multiple measures from the same bird. To investigate whether bird diet treatment influenced change in consumption time across manipulation days, we also tested for an interaction between the diet treatment and the manipulation day.

The effect of bird diet treatment on body weight was analyzed by calculating the difference between a bird’s capture weight and the weight after the six day diet manipulation (g), and analysing this difference using a linear model with bird diet treatment as a fixed effect (n = 40). The models for physiological markers were run using the mean values of the duplicates or triplicates, except for the mitochondrial respiration measures that were analyzed only once. If the assays were done using several plates (corticosterone, TBARS, GSH, GST), we controlled inter-assay differences by including plate number as a fixed effect and retaining this in the model if it was statistically significant. To test the hypothesis that PAs negatively affected the oxidative status of great tits, we performed a separate model for each oxidative stress biomarker (TBARS, GST, GSH, GSH:GSSG). The final models for GSH:GSSG ratio (ln-transformed, n = 60) and lipid peroxidation (TBARS, n = 59) included bird diet treatment (C no NF/ C + NF/ PA + no NF/ PA + NF) and assay plate number as fixed effects, and the final models for GSH concentration (ln-transformed, n = 63) and GST activity (ln-transformed, n = 41) included only bird diet treatment. The final model for corticosterone concentration (log_10_-transformed, n = 59) included bird diet treatment and plate number. When analysing serotonin concentration (log_10_-transformed, n = 37), the final model also included body condition. Regarding the mitochondrial respiration metrics, the final models for endogenous respiration (ROUTINE) and flux control ratios (FCR ROUTINE/(CI+II)) included bird diet treatment, sex and cell count, and the final model for oxidative phosphorylation (OXPHOS) included bird diet treatment and age (n = 43 in all models).

All statistical analyses were conducted using R version 4.4.3 (R Core Team, 2025). Linear models were done using *lme4* (Bates et al. 2015) and *lmerTest* packages (Kuznetsova et al. 2017). Normality and heteroscedasticity of residuals were checked with the *DHARMa* package (Hartig et al. 2024). The significance of the main effect of bird diet treatment was tested using the Anova function in the *car* package (Fox and Weisberg 2019). If the main effect of treatment was significant, pairwise post-hoc comparisons were performed using the *emmeans* package (Lenth 2025). Graphs were created using the *ggplot2* package (Wickham 2016).

## RESULTS

### Time to consume the moths

The bird diet treatment had a significant effect on the time to consume the moth pellet (χ² = 17.073, *p* < 0.001; Fig. 2). Birds in the C no NF treatment (lowest PA concentration) consumed the moth pellets more quickly compared to the two treatments where birds received moths reared on the PA diet (compared to PA no NF: estimate = −47.088 ± 15.304, *t* = −3.077, *p* = 0.016; compared to PA + NF: estimate = −54.987 ± 15.518, *t* = −3.543, *p* = 0.004). There was a tendency for birds in the C + NF treatment to consume the moths more quickly than birds in the PA + NF treatment (estimate = −39.949 ± 15.538, *t* = −2.571, *p* = 0.059), but this difference was not observed between C + NF and PA no NF treatments (estimate = −32.051 ± 15.324, *t* = −2.092, *p* = 0.17). There was no significant difference in the time to consume the moths between the two control (C no NF vs. C + NF: estimate = −15.038 ± 15.497, *t* = −0.970, *p* = 0.77), or the two PA treatments (PA no NF vs. PA + NF: estimate = −7.898 ± 15.345, *t* = −0.515, *p* = 0.96). Regardless of the treatment (manipulation day × treatment: χ² = 5.720, *p* = 0.13), birds consumed the moths more quickly as the experiment progressed (effect of manipulation day: estimate = −11.484 ± 2.195, *t* = −5.232, *p* < 0.001; Fig. 2). A separate model comparing birds that consumed moths reared on PA and control diets showed a similar result, with birds receiving control moths consuming the pellets significantly more quickly compared to the birds receiving PA moths (estimate = −43.426 ± 10.840, *t* = −4.006, *p* < 0.001; Fig. 2).

**Figure 2.**
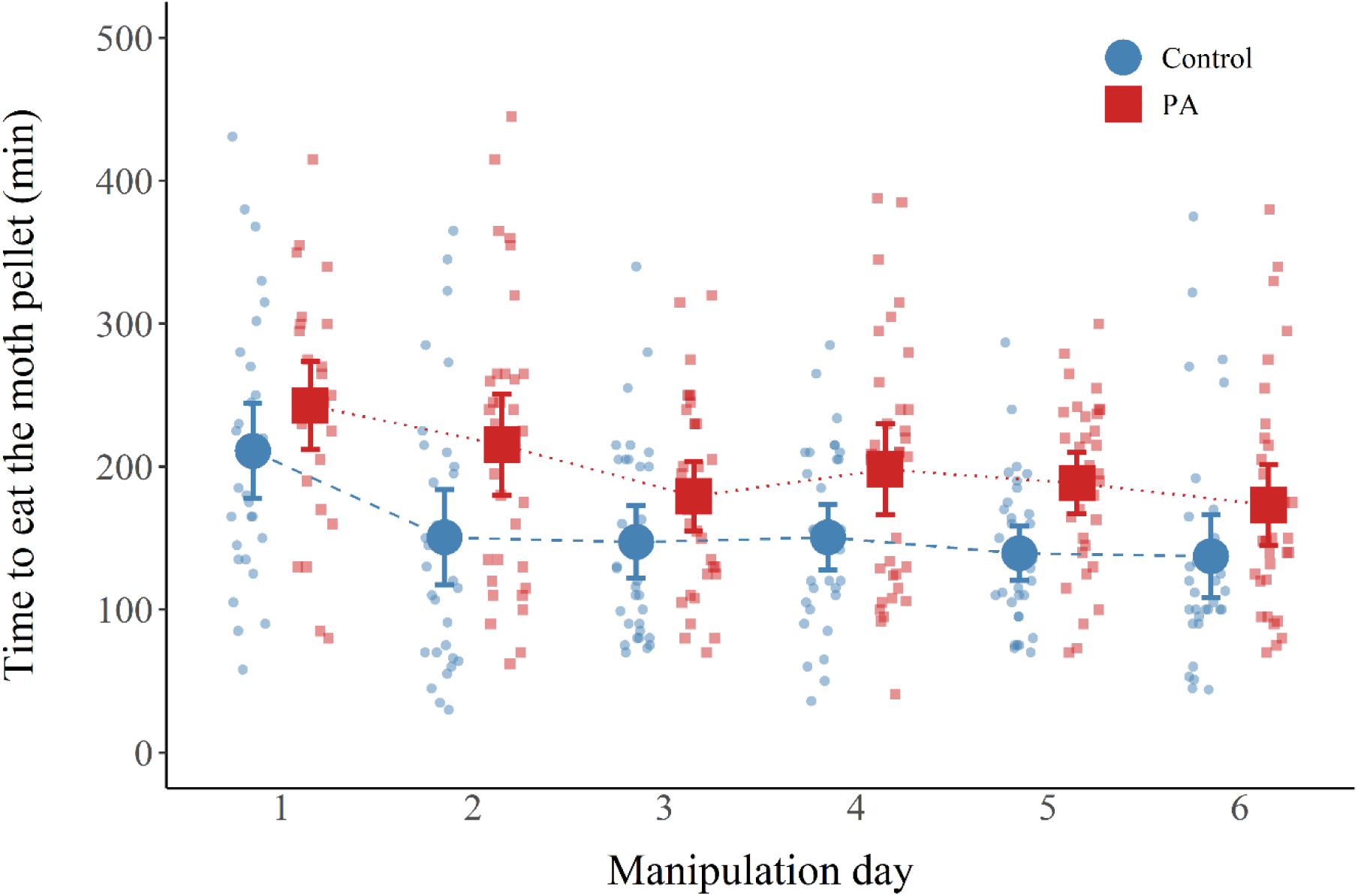
Time (min) taken to eat the moth pellet across the six diet manipulation days for birds that received moths reared on the control (blue circles) or on the PA diet (red squares). The colored symbols show the mean and the vertical error bars indicate 95% confidence intervals. Smaller symbols indicate individual variation within the treatment.

### Body weight and corticosterone concentration

Regardless of the diet treatment, the weight of the birds decreased during the experiment (mean weight loss = 1.5g), which is typical when wild birds are brought into captivity. There was no difference in weight change among the four bird diet treatments (*F*_3,36_ = 0.300, *p* = 0.82), or when comparing birds that received moths reared on control and PA diets (estimate = 0.146 ± 0.256, *t* = 0.570, *p* = 0.57). We also found no effect of bird diet treatment on their corticosterone levels (*F*_3,54_ = 0.264, *p* = 0.85; Fig. 3). Likewise, there were no significant differences in corticosterone levels when we compared birds that consumed moths reared on control and PA diets (estimate = −0.019 ± 0.048, t = −0.403, *p* = 0.69).

**Figure 3.**
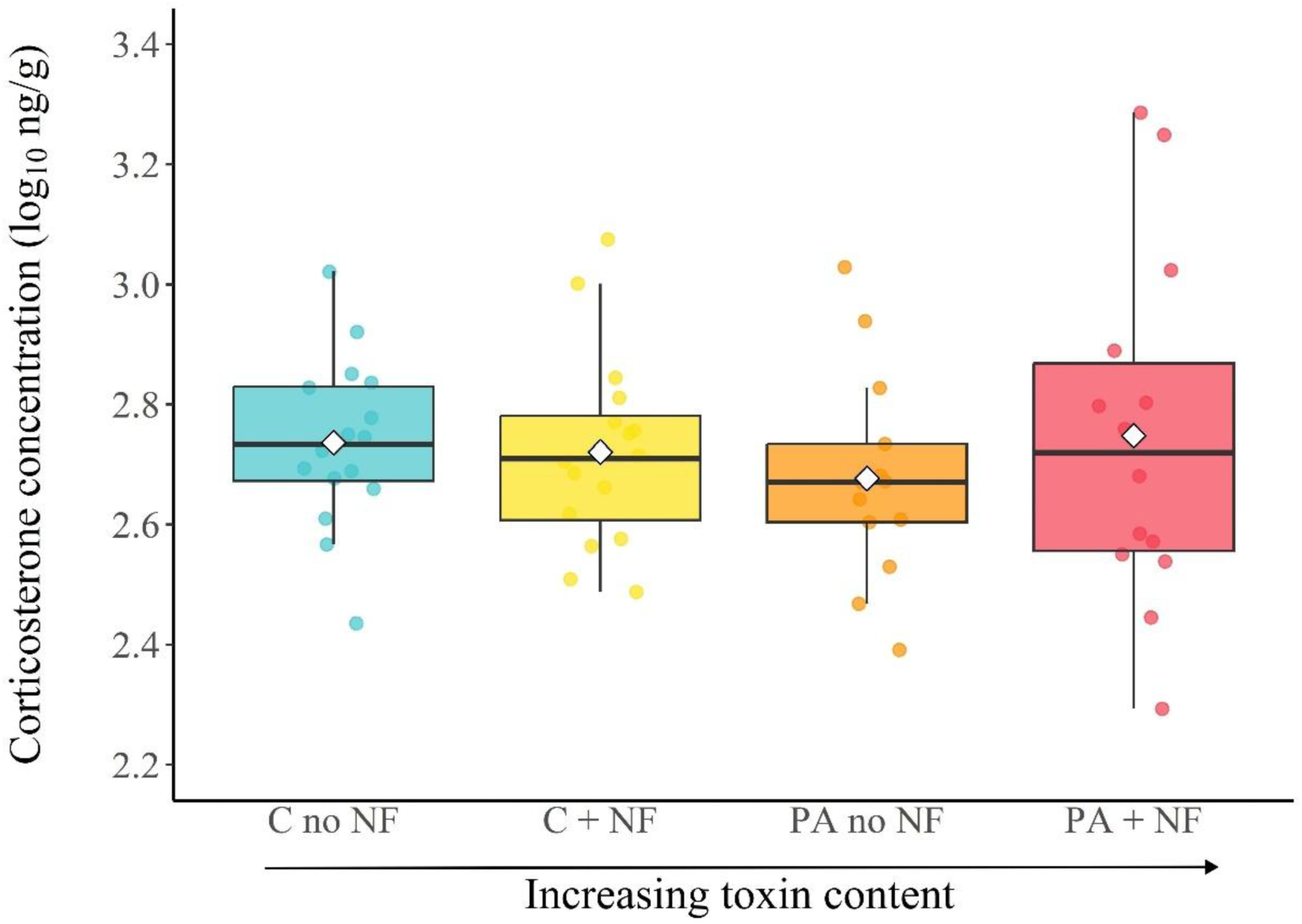
Corticosterone concentrations (ng/g, log_10_-transformed) in each bird diet treatment. Boxplots represent the median (horizontal line within the box), the first and third quartiles (lower and upper bounds of the box), and the whiskers indicate the values within 1.5 times the interquartile range. Circles represent individual values for each sample, while white diamonds indicate the group mean.

### Oxidative stress

The consumption of the moths of different levels of toxicity did not influence any of the analyzed biomarkers of oxidative stress or damage, with no significant effect of bird diet treatment on GSH concentration (*F*_3,59_ = 0.114, *p* = 0.95), GSH:GSSG ratio (*F*_3,55_ = 0.792, *p* = 0.50), the activity of GST (*F*_3,37_ = 0.825, *p* = 0.49), or lipid peroxidation (*F*_3,53_ = 2.169, *p* = 0.10; Fig. 4). Similarly, we found no significant differences between birds that consumed moths reared on control and PA diets in GSH concentration (estimate = 0.031 ± 0.083, *t* = 0.373, *p* = 0.71), in GSH:GSSG ratio (estimate = −0.085 ± 0.127, *t* = −0.674, *p* = 0.50), in the activity of GST (estimate = 0.147 ± 0.155, t = 0.953, *p* = 0.35), or in lipid peroxidation (estimate = 0.001 ± 0.001, *t* = 1.629, *p* = 0.11).

**Figure 4.**
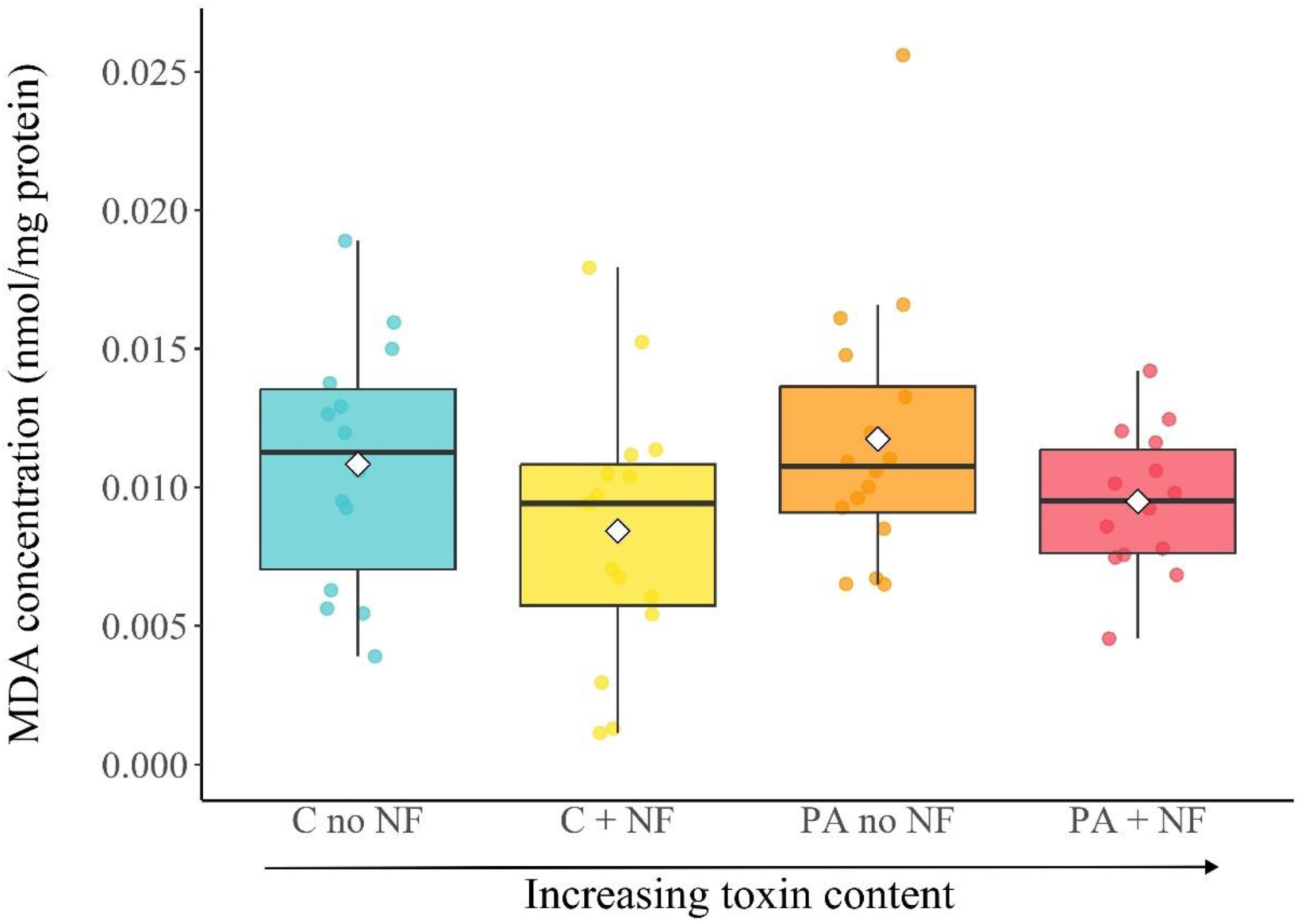
Lipid peroxidation (malondialdehyde, MDA, concentration, nmol/mg of protein) in each bird diet treatment. Boxplots represent the median (horizontal line within the box), the first and third quartiles (lower and upper bounds of the box), and the whiskers indicate the values within 1.5 times the interquartile range. Circles represent individual variation and white diamonds indicate the mean for each treatment.

### Mitochondrial metabolism

There was a tendency for birds receiving moths with lowest PA concentration (C no NF) to have lower endogenous respiration (ROUTINE) than other treatments (see Fig. 5a and appendix Table A10a), but the main effect of treatment was not statistically significant (*F*_3,37_ = 2.305, *p* = 0.093). There was no difference in endogenous respiration between males and females (estimate = 0.048 ± 0.030 *t* = 1.632, *p* = 0.11), but sex was kept in the final model based on AIC values (see appendix). As expected, cell count had a significant positive effect on endogenous respiration, with respiration rates increasing with increasing cell numbers (estimate = 0.031 ± 0.011 *t* = 2.742, *p* = 0.009). We found no effect of bird diet treatment on respiration through oxidative phosphorylation (OXPHOS, *F*_3,38_ = 0.429, *p* = 0.73; Fig. 5b). Age was kept in the final model based on AIC values even though it did not have a significant effect on OXPHOS (estimate = 0.007 ± 0.004 *t* = 1.593, *p* = 0.12). Finally, we found no significant differences between birds that consumed moths reared on control and PA diets on endogenous respiration (estimate = 0.036 ± 0.031, *t* = 1.166, *p* = 0.25), or on OXPHOS (estimate = −0.0009 ± 0.004, *t* = −0.240, *p* = 0.81).

**Figure 5.**
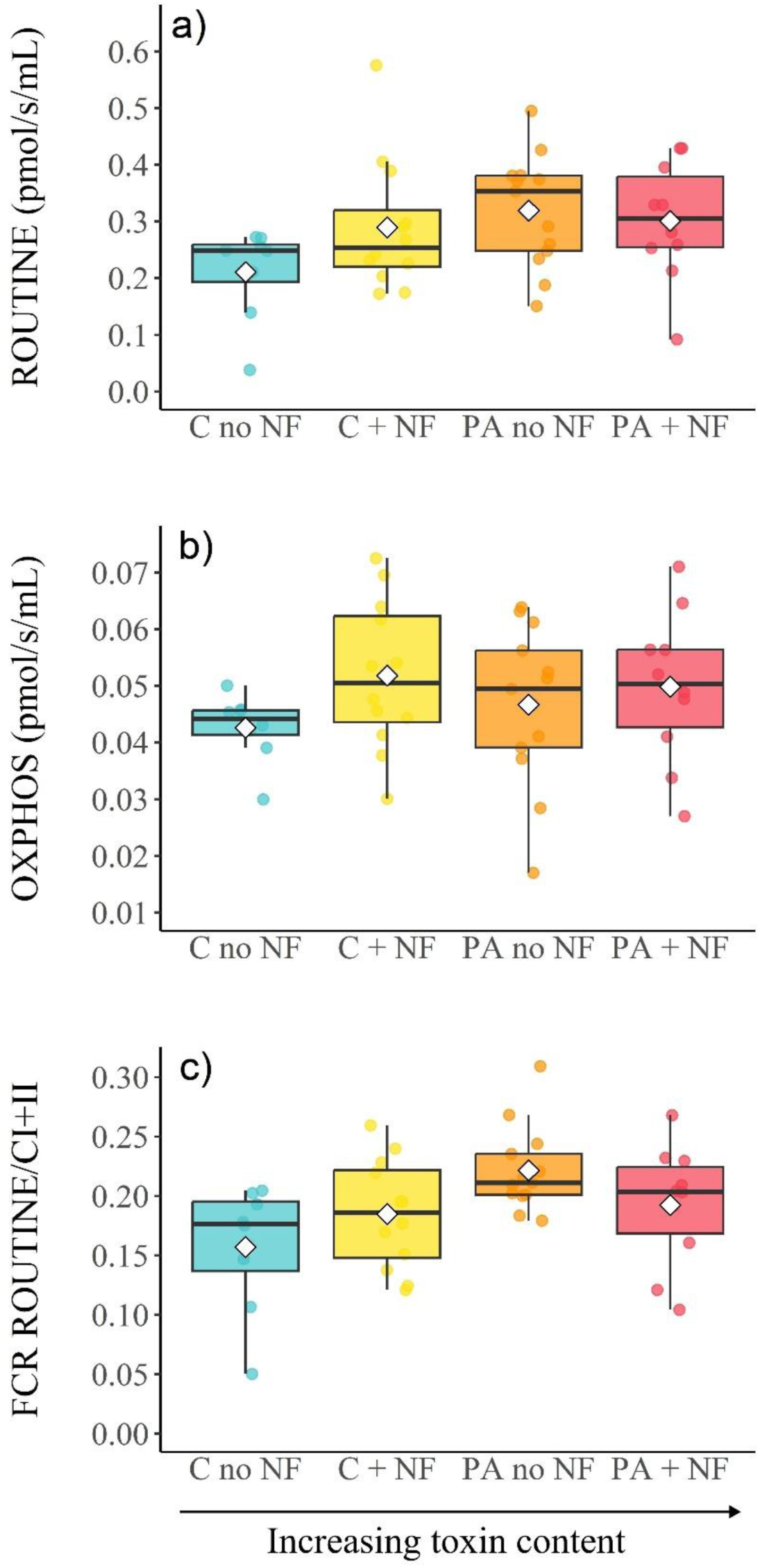
Mitochondrial metabolism in different treatment groups, including a) routine respiration (pmol/s/mL), b) OXPHOS (pmol/s/mL), and c) FCR ROUTINE/CI+CII ratio. Boxplots represent the median (horizontal line within the box), the first and third quartiles (lower and upper bounds of the box), and the whiskers indicate the values within 1.5 times the interquartile range. Circles represent individual variation and white diamonds indicate the mean for each treatment.

In contrast, bird diet treatment significantly influenced the usage of mitochondrial maximal respiration capacity (FCR ROUTINE/(CI+II), *F*_3,37_ = 5.013, *p* = 0.005; Fig. 5c). Post-hoc test comparisons among the four treatments showed that birds in the PA no NF treatment exhibited significantly higher FCR values than those in the C no NF group (estimate = 0.069 ± 0.018, *t* = 3.729, *p* = 0.003, Fig. 5c), but there were no significant differences among other treatments (all other post-hoc comparisons *p* > 0.05). We also found an effect of treatment when comparing birds that consumed moths reared on control and PA diets, with birds that received PA moths having higher FCR values (estimate = 0.031 ± 0.014, *t* = 2.222, *p* = 0.03). Finally, we found that a bird’s sex had an effect on FCR values, with males showing higher FCR values than females (estimate = 0.042 ± 0.013, *t* = 3.271, *p* = 0.002).

### Serotonin concentration

Plasma serotonin concentrations did not differ significantly among bird diet treatments groups (*F*_3,32_ = 0.085, *p* = 0.97, Fig. 6). There was a tendency, although not quite statistically significant, for birds with higher body condition to have higher serotonin levels (estimate = 0.118 ± 0.066, *t* =1.801, *p* = 0.081). When comparing birds that consumed moths reared on control and PA diets, there was no significant difference in serotonin concentration (estimate = 0.015 ± 0.148, *t* = 0.103, *p* = 0.92).

**Figure 6:**
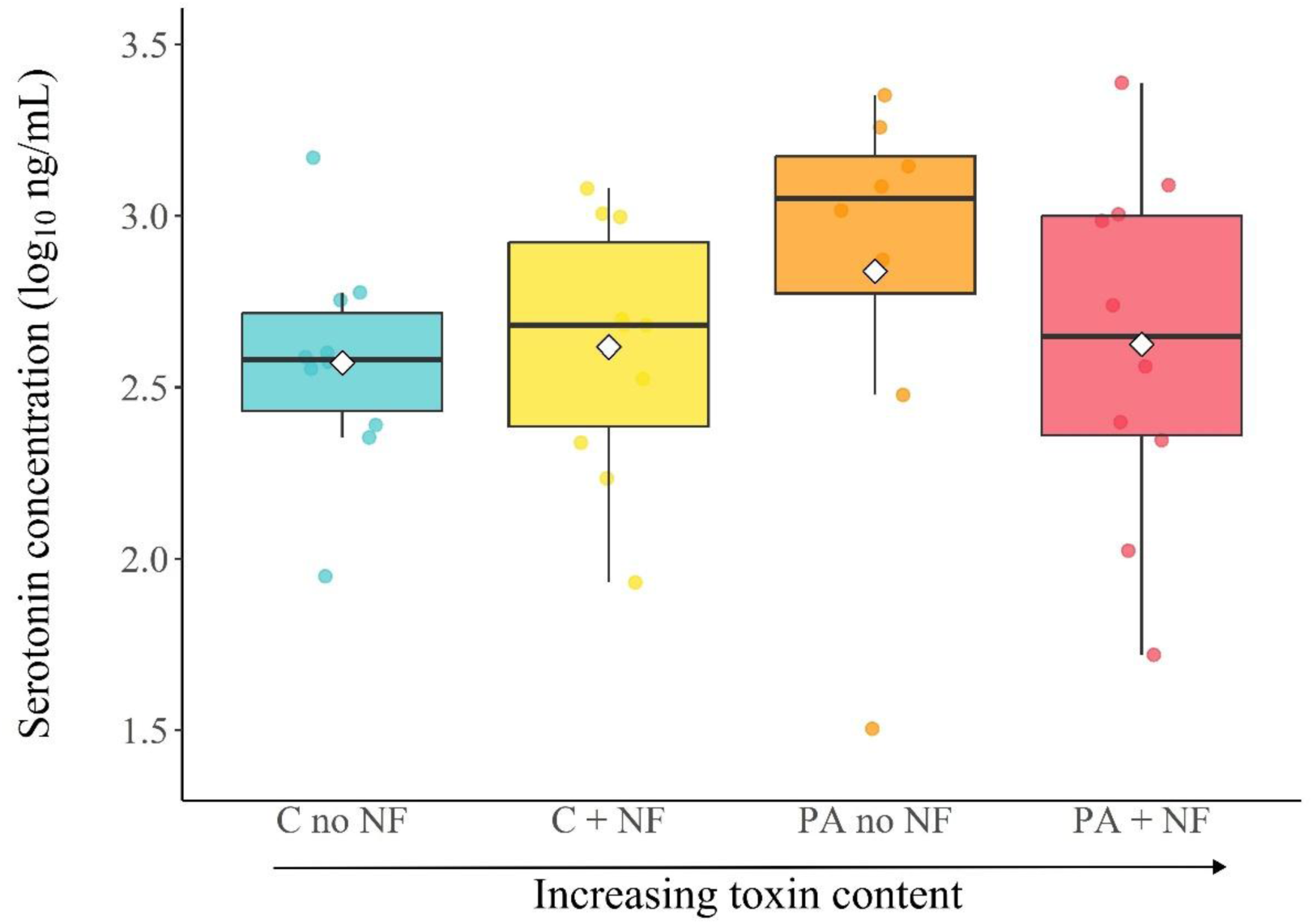
Plasma serotonin concentrations (ng/mL, log_10_-transformed) for the different treatment groups. Boxplots represent the median (horizontal line within the box), the first and third quartiles (lower and upper bounds of the box), and the whiskers indicate the values within 1.5 times the interquartile range. Circles represent individual values for each sample, while white diamonds indicate the group mean.

## DISCUSSION

In this study, we explored the physiological effects of the consumption of chemically defended insect prey on an avian predator. Many insects sequester toxic pyrrolizidine alkaloids (PAs) as a defence against predators (Rothschild et al. 1979; Von Nickisch-Rosenegk and Wink 1993; Trigo 2011). We predicted that great tits receiving moths with higher PA concentrations would exhibit elevated corticosterone levels and oxidative stress, along with reduced serotonin concentrations, but we did not find significant differences among bird diet treatments in any of these biomarkers. Regarding mitochondrial metabolism, we found that the usage of mitochondrial maximal respiration capacity was higher for the birds that received moths with higher PA concentrations, but this effect was moderate and needs to be interpreted with caution due to our limited sample size. Altogether, our results suggest that a low consumption of chemically defended prey does not induce significant physiological costs to avian predators.

Our experiment included four diet treatments where great tits received *A. plantaginis* moths with varying PA profiles. The main PAs identified in *A. plantaginis* include senecionine and seneciphylline (Winters et al. 2021), and we quantified both their free base forms, as well as less toxic N-oxides. The N-oxides are suggested to be the storage form of PAs in moths (Opitz and Müller 2009), and we found that their concentration was considerably higher in the moths raised on the PA diet. Interestingly, free base seneciphylline and senecionine concentrations decreased significantly when the defensive neck fluids of the moths were removed, suggesting that the more toxic forms of PAs are stored in the defensive neck fluids. The neck fluids have been previously demonstrated to contain methoxypyrazines (Burdfield-Steel et al. 2018), which act as an olfactory deterrent and can interact with PAs to influence predator attack decisions (Winters et al. 2021). We did not quantify methoxypyrazines in our experiment because they are highly volatile (Khashaveh et al. 2025), and it is unlikely that any significant amounts remained after the freeze-drying process. Surprisingly, we found low concentrations of PAs also in the moths reared in the control diet that consisted of dandelion only (Table 1), which to our knowledge has not been reported to contain PAs. Many arctiids incorporate PAs into their eggs (Dussourd et al. 1988; Opitz and Müller 2009), and because we collected the females from the wild without controlling their diet, the larvae might have been able to accumulate some PAs from the eggs. It is also possible that dandelions contain small amounts of PAs and the moths are very efficient at sequestering these. Nevertheless, the concentrations detected in the control treatment were considerably lower than those observed in the PA treatment, so it is unlikely that the lack of a treatment without any PAs influenced our conclusions.

The PA concentrations used in this experiment may have been insufficient to bring out detectable changes in the measured physiological biomarkers. However, our aim was to use doses that were assumed to be ecologically relevant. While it is known that predators can make strategic decisions to include toxic prey in their diet (Skelhorn and Rowe 2007; Barnett et al. 2007, 2012), accurately estimating the natural toxin intake in wild birds is challenging because predation events of individual birds are often difficult to observe. Furthermore, predators do not necessarily always consume the entire prey, as they can learn to selectively ingest the least toxic body parts (Brown and Neto 1976; Fink et al. 1981), which makes determining the toxin load even more challenging. Our finding of increased latency for birds to consume the moth pellets with high PA concentrations indicates that birds perceived these moths more unpalatable, possibly because of their bitter taste (Glendinning 1994). We did not observe birds’ other behaviors during the diet manipulation, but previous studies suggest that PAs elicit aversive responses (e.g. beak wiping and head shaking) and prey rejection in avian predators (Rowell-Rahier et al. 1995; Cardoso 1997). Including higher toxin concentrations in our study would have been therefore challenging, as birds might have refused to consume the prey. In addition to not being ecologically relevant, the use of very high toxin concentrations would also have ethical implications.

To our knowledge, this is the first study that has quantified the physiological effects of insect PA defences on wild avian predators. Previous work on PA toxicity on birds has focused on domestic species, such as chickens and quails, feeding on PA-containing plants or their seeds (Buckmaster et al. 1977; Eröksüz et al. 2003, 2008; Bezerra et al. 2011; Diaz et al. 2014; Brown et al. 2016). Typical adverse effects observed in these studies include negative effects on growth and egg production, and different degrees of liver damage and changes in associated liver enzymes (e.g. Eröksüz et al. 2003, 2008; Diaz et al. 2014; Brown et al. 2016). However, the used alkaloids, their concentrations and exposure times vary greatly among studies, which makes the results hard to compare. In our study, the highest toxin treatment (PA + NF) included a daily dose of approximately 0.75mg/kg Senecionine and 0.5mg/kg Seneciphylline, but the moths also contained the corresponding N-oxides that might be reduced back into the more toxic free base forms in the gut of the predators (Nuringtyas et al. 2014; Yang et al. 2019). Furthermore, we only quantified the previously identified compounds in *A. plantaginis* (Winters et al. 2021), but the moths might also have other PAs, with over 660 types of PAs identified in nature (Xu et al. 2019), so the total PA concentration in our study might have been higher than we estimated. Previous studies have demonstrated that even low doses of PAs can have adverse impacts. For example, a five-week exposure to a daily dose of 0.54mg/kg usaramine from *Crotalaria pallida* seeds decreased body weight and increased enzymes associated with liver damage in hens (Diaz et al. 2014) and a seven-day exposure of 5mg/kg *C. retusa* seeds caused liver damage in guinea fowl (*Numida meleagris*, Bezerra et al. 2011). However, species vary highly in their susceptibility to PAs (White et al. 1973; He et al. 2021). From the tested avian species, quails seem to be particularly resistant (Buckmaster et al. 1977; Eröksüz et al. 2008). For example, LD_50_ for retrorsine was found to be 279mg/kg in quail, compared to 85mg/kg in fowl and 34mg/kg in rats (White et al. 1973). This variation among species, as well as the observed toxicity differences of different alkaloids and their combinations (Brown et al. 2016), highlights the importance of studying the physiological costs of prey toxins using ecologically relevant predators and prey.

In contrast to previous studies that have often found PA consumption decreasing body weight (Eröksüz et al. 2008; Diaz et al. 2014; Brown et al. 2016), we did not see this trend after the diet manipulation. This could be explained by a short exposure time, with a previous study in hens similarly finding no differences after one week exposure to usaramine but a significant decrease in weight after two weeks (Diaz et al. 2014). Against our prediction, we also did not detect any differences in oxidative stress, or corticosterone or serotonin concentrations among diet treatments. While this result might be explained by great tits being resistant to low PA doses, it is also possible that the biomarkers that we measured did not capture the toxicity effects or the effects are tissue-specific. For example, previous studies that have reported PAs to induce oxidative stress have often quantified changes in hepatic GSH or GST activity (Xu et al. 2019) rather than blood oxidative stress biomarkers used in our study. Hepatic sampling was not feasible in our study due to ethical constraints of working with wild birds. In addition, our results could potentially be influenced by captive conditions, which are known to cause stress to wild birds (Fischer and Romero 2019) and might make it harder to detect any PA-induced changes. For example, newly captured birds often lose weight (Fischer and Romero 2019), which was also observed in our study, with the weight of the birds decreasing during the six-day manipulation regardless of the diet treatment. Finally, while the captive conditions in our study were standardized, in nature the consumption of prey toxins could be associated with other stressors, such as low food availability that is predicted to increase predator attacks on aposematic prey (Sherratt 2003; Barnett et al. 2007). These other factors could potentially influence toxin resistance, but exploring these interactive effects in wild birds is challenging.

The only physiological changes detected in our study were related to mitochondrial metabolism. There was a tendency for routine respiration to increase when the consumed PA dose increased, although this effect was not statistically significant. We also found that the coupling efficiency ratio of ROUTINE/(CI+II) increased when birds received moths with higher PA concentrations, which suggests that the cells operated closer to their maximum respiratory capacity. This supports the hypothesis that elevated PA exposure induces metabolic stress, leading to a compensatory increase in mitochondrial activity, and potentially reflecting a state of mitochondrial stress and reduced energetic efficiency. We did not find significant differences among treatments in the OXPHOS ratio, suggesting that ATP production efficiency remained unchanged but greater respiratory effort was required to maintain ATP output. A previous study demonstrated the effect of PAs on hepatic mitochondrial metabolism in rats, showing that reactive pyrrole-protein adducts can bind to ATP synthase subunit beta (ATP5B), leading to reduced ATP synthase activity and intracellular ATP levels, as well as reduced mitochondrial membrane potential and respiration (Lu et al. 2018). Our study suggests that PAs might also influence blood cell mitochondrial metabolism, although the detected changes were moderate, and further studies with higher PA doses and other tissues are required to get a better understanding of these effects. Finally, we found that males used more of their mitochondrial maximum capacity than females. This is similar to a previous study with another passerine species (pied flycatchers, *Ficedula hypoleuca*), although this was done in a reproductive context (Stier et al. 2019), whereas our study was conducted outside the breeding season.

While there has been extensive theoretical (Sherratt 2003) and empirical work (Barnett et al. 2007, 2012; Skelhorn and Rowe 2007) examining predator decision making and trade-offs of attacking aposematic prey, the physiological costs of ingesting prey toxins have remained largely unknown. PAs are common chemical defences found in many insects (Trigo 2011), but most previous studies on their toxicity have focused on food safety, human health and poisoning in domestic animals (Wiedenfeld and Edgar 2011). Here, we have shown that a low consumption of moths with PA defences (one defended moth per day) does not have significant physiological costs to great tit predators, at least in the biomarkers and tissues we analyzed. One limitation in our study is a small sample size in some of the analyses, which might have reduced statistical power to detect differences among treatments. In future studies, it would be interesting to obtain multiple blood and fecal samples throughout the diet manipulation to follow the physiological changes immediately after the toxin exposure. While we focused here on PA defences, future work should also aim to quantify the costs of different types of prey toxins and their potentially interactive effects, as wild predators are likely to consume different aposematic insects that might contain various combinations of defensive compounds. This includes investigating potential long-term fitness effects on predator survival or reproduction, in addition to immediate physiological changes. Overall, our study opens up new research avenues to start unravelling the links between physiological costs of prey toxin consumption, predator foraging decisions and predator-prey coevolution.

## ACKNOWLEDGEMENTS

We thank Johanna Mappes for the discussions and advice on moth rearing, Ella Ruutu for the help with the experiment, and the staff at the Konnevesi research station for providing facilities for the birds. The project was funded by the Academy of Finland Research Fellowship (#355869) and a Horizon Europe MSCA Postdoctoral Fellowship (#101151042) awarded to LH.

## Notes

### Competing Interest Statement

The authors have declared no competing interest.

## LITERATURE CITED

Bates D., M. Mächler, B. Bolker, and S. Walker. 2015. Fitting linear mixed-effects models using lme4. J Stats Soft 67:1–48.

Barnett C.A., J. Skelhorn, M. Bateson, and C. Rowe. 2012. Educated predators make strategic decisions to eat defended prey according to their toxin content. Behav Ecol 23:418–424.

Barnett C.A., M. Bateson, and C. Rowe. 2007. State-dependent decision making: educated predators strategically trade off the costs and benefits of consuming aposematic prey. Behav Ecol 18:645–651.

Bezerra W.G.A., M.A. de Souza, R. Horn, L.A.V.C. Vasconcelos, J.S. Batista, and B. Soto-Blanco. 2011. Pathological effects of short-term *Crotalaria retusa* ingestion by guinea fowl (*Numida meleagris*). IJPPR 1:65–69.

Brown A.W., B.L. Stegelmeier, S.M. Colegate, D.R. Gardner, K.E. Panter, E.L. Knoppel, and J.O. Hall. 2016. The comparative toxicity of a reduced, crude comfrey (*Symphytum officinale*) alkaloid extract and the pure, comfrey-derived pyrrolizidine alkaloids, lycopsamine and intermedine in chicks (*Gallus gallus domesticus*). J Appl Toxicol 36:716–725.

Brown Jr, K.S. and J.V. Neto. 1976. Predation on aposematic ithomiine butterflies by tanagers (*Pipraeidea melanonota*). Biotropica 8:136–141.

Buckmaster G.W., P. R. Cheeke, G.H. Arscott, E.O. Dickinson, M.L. Pierson, and L.R. Shull. 1977. Response of Japanese quail to dietary and injected pyrrolizidine (*Senecio*) alkaloid. J Anim Sci 45:1322–1325.

Burdfield-Steel E., H. Pakkanen, B. Rojas, J.A. Galarza, and J. Mappes. 2018. De novo synthesis of chemical defenses in an aposematic moth. J Insect Sci. 18:1–4.

Cardoso M.Z. 1997. Testing chemical defence based on pyrrolizidine alkaloids. Anim Behav 54:985–991.

Cardoso S.M., C. Pereira, and C.R. Oliveira. 1999. Mitochondrial function is differentially affected upon oxidative stress. Free Radic Biol Med 26:3–13.

Cheeke P.R. and M.L. Pierson-Goeger. 1983. Toxicity of *Senecio jacobaea* and pyrrolizidine alkaloids in various laboratory animals and avian species. Toxicol Lett 18:343–349.

Cheng D., V.T. Nguyen, N. Ndihokubwayo, J. Ge, and P.P.J. Mulder. 2017. Pyrrolizidine alkaloid variation in *Senecio vulgaris* populations fro m native and invasive ranges. PeerJ 5:e3686.

Costantini D. 2008. Oxidative stress in ecology and evolution: Lessons from avian studies. Ecol Lett 11:1238–1251.

Diaz G.J., L.X. Almeida, and D.R. Gardner. 2014. Effects of dietary *Crotalaria pallida* seeds on the health and performance of laying hens and evaluation of residues in eggs. Res Vet Sci 97:297–303.

Dussourd D.E., K. Ubik, C. Harvis, J. Resch, J. Meinwald, and T. Eisner. 1988. Biparental defensive endowment of eggs with acquired plant alkaloid in the moth *Utetheisa ornatrix*. Proc Natl Acad Sci 85:5992–5996.

Eröksüz H., Y. Eröksüz, H. Özer, I. Yaman, F. Tosun, C. Akyüz Kizilay, and U. Tamer. 2003. Toxicity of Senecio vernalis to laying hens and evaluation of residues in eggs. Vet Hum Toxicol 45:76–80.

Eröksüz Y., A.O. Çeribaşi, A. Cevik, H. Eröksüz, F. Tosun, and U. Tamer. 2008. Toxicity of *Heliotropium dolosum*, *Heliotropium circinatum*, and *Senecio vernalis* in parental quail and their progeny, with residue evaluation of eggs. Turk J Vet Anim Sci 32:475–482.

Espín S., P. Sánchez-Virosta, and A.J. García-Fernández. 2017. A microplate adaptation of the thiobarbituric acid reactive substances assay to determine lipid peroxidation fluorometrically in small sample volumes. Rev Toxicol 34:94–98.

Fink L.S. and L.P. Brower. 1981. Birds can overcome the cardenolide defence of monarch butterflies in Mexico. Nature 291:67–70.

Fischer C.P. and L.M. Romero. 2019. Chronic captivity stress in wild animals is highly species-specific. Conserv Physiol 7, coz093.

Fox J. and S. Weisberg 2019. An R Companion to applied regression, 3rd ed. Sage, Thousand Oaks, CA.

Fu P.P., Q. Xia, G. Lin, and M.W. Chou. 2004. Pyrrolizidine alkaloids—genotoxicity, metabolism enzymes, metabolic activation, and mechanisms. Drug Metab Rev 36:1–55.

Glendinning J.I. 1994. Is the bitter rejection response always adaptive? Physiol Behav 56:1217–1227.

Gordon S.P., E. Burdfield-Steel, J. Kirvesoja, and J. Mappes. 2021. Safety in numbers: How color morph frequency affects predation risk in an aposematic moth. Am Nat 198:128–141.

Halliwell B. and J.M.C. Gutteridge. 2015. Free Radicals in Biology and Medicine. 5th ed. Oxford University Press, New York.

Hartig F. 2024. DHARMa: Residual diagnostics for hierarchical (multi-level / mixed) regression models. R package version 0.4.7.

Harvey P.W., G. Healing, S.J. Rees, D.J. Everett, and A. Cockburn. 1994. Glucocorticosteroid interactions with natural toxins: a mini review. Nat Toxins 2:341–346.

He Y., L. Zhu, J. Ma, and G. Lin. 2021. Metabolism-mediated cytotoxicity and genotoxicity of pyrrolizidine alkaloids. Arch Toxicol 95:1917–1942.

Hess L. 2002. Corticosteroid synthesis and metabolism in birds. Semin Avian Exot Pet Med 11:65–70.

Isaksson C., J. Örnborg, E. Stephensen, and S. Andersson. 2005. Plasma glutathione and carotenoid coloration as potential biomarkers of environmental stress in great tits. EcoHealth 2:138–146.

Khashaveh A., C. Yi, H. Tang, X. Song, G. Zhang, J. Xie, and Y. Zhang. 2025. Biological roles of pyrazines in insect chemical communication. Chem Biol Technol Agric 12:109.

Kuznetsova A., P.B. Brockhoff, and R.H.B. Christensen. 2017. lmerTest Package: Tests in linear mixed effects models. J Stat Soft 82:1–26.

Leach J., H.N. Suber, K. Conley, R. Rivera, J. Surles, B. Hames, and R.J. Kendall. 2024. Validation of an enzyme immunoassay for the detection of corticosterone metabolites from northern bobwhite (*Colinus virginianus*) feces. Conserv Physiol 12, coad098.

Lenth R. 2025. emmeans: Estimated marginal means, aka least-squares means. R package version 1.11.0.

Liang Q.N., T. Liu, L. Ji, Y. Min, and Y. Xia. 2009. Pyrrolizidine alkaloid clivorine-induced oxidative stress injury in human normal liver L-02 cells. Drug Discov Ther 3:247–251.

Lu Y., J. Ma, Z. Song, Y. Ye, P.P. Fu, and G. Lin. 2018. The role of formation of pyrrole–ATP synthase subunit beta adduct in pyrrolizidine alkaloid-induced hepatotoxicity. Arch Toxicol 92:3403–3414.

Manoli I., S. Alesci, M.R. Blackman, Y.A Su, O.M. Rennert, and G.P. Chrousos. 2007. Mitochondria as key components of the stress response. Trends Endocrinol Metab 18:190–198.

Mingatto F.E., D.J. Dorta, A.B. dos Santos, I. Carvalho, C.H.T.P da Silva, V.B. da Silva, S.A. Uyemura, A.C. dos Santos, and C. Curti. 2007. Dehydromonocrotaline inhibits mitochondrial complex I. A potential mechanism accounting for hepatotoxicity of monocrotaline. Toxicon 50:724–730.

Mohammad-Zadeh L.F., L. Moses, and S.M. Gwaltney-Brant. 2008. Serotonin: a review. J Vet Pharmacol Ther 31:187–199.

Nokelainen O., J. Valkonen, C. Lindstedt, and J. Mappes. 2014. Changes in predator community structure shifts the efficacy of two warning signals in Arctiid moths. J Anim Ecol 83:598–605.

Nuringtyas T.R., R. Verpoorte, P.G. Klinkhamer, M.M. van Oers, and K.A. Leiss. 2014. Toxicity of pyrrolizidine alkaloids to *Spodoptera exigua* using insect cell lines and injection bioassays. J Chem Ecol 40:609–616.

Opitz S.E. and C. Müller. 2009. Plant chemistry and insect sequestration. Chemoecology 19:117–154.

Peig J. and A. Green. 2009. New perspectives for estimating body condition from mass/length data: The scaled mass index as an alternative method. Oikos 118:1883–1891.

Poulton S.E.B. 1890. The Colours of Animals: Their Meaning and Use, Especially Considered in the Case of Insects. Kegan Paul, Trench, Trübner & Co, London.

R Core Team. 2025. R: A language and environment for statistical computing. R Foundation for Statistical Computing, Vienna. http://www.R-project.org/.

Rainio M.J., T. Eeva, T. Lilley, J. Stauffer, and S. Ruuskanen. 2015. Effects of early-life lead exposure on oxidative status and phagocytosis activity in great tits (*Parus major)*. Comp Biochem Physiol C Toxicol Pharmacol 167:24–34.

Reinwaldt C., M. Bittner, G. Kempe, and T. Glauner. 2017. Quantitation of pyrrolizidine alkaloids in honey and herbal teas by UHPLC/MS/MS. Agilent Technologies Incorporation.

Rothschild M., R.T. Aplin, P.A. Cockrum, J.A. Edgar, P. Fairweather, and R. Lees. 1979. Pyrrolizidine alkaloids in arctiid moths (Lep.) with a discussion on host plant relationships and the role of these secondary plant substances in the Arctiidae. Biol J Linn Soc 12:305–326.

Rowell-Rahier M., J.M. Pasteels, A. Alonso-Media, and L.P. Brower. 1995. Relative unpalatability of leaf beetles with either biosynthesized or sequestered chemical defence. Anim Behav 49:709–714.

Royama T. 1970. Factors governing the hunting behaviour and selection of food by the great tit (*Parus major* L.). J Anim Ecol 39:619–668.

Ruxton G.D., W.L. Allen, T.N. Sherratt, and M.P. Speed. 2018. Avoiding attack: the evolutionary ecology of crypsis, aposematism, and mimicry. 2nd ed. Oxford University Press, Oxford.

Saldanha B.C., P.A. Silva, C. Maximino, G.C. Cardoso, S. Trigo, and M.C. Soares. 2023. The role of serotonin in modulating common waxbill behaviour. Behav Ecol Sociobiol 77:39.

Sapolsky R.M., L.M. Romero, and A.U. Munck. 2000. How do glucocorticoids influence stress responses? Integrating permissive, suppressive, stimulatory, and preparative actions. Endocr Rev 21:55–89.

Schmeller T., A. El-Shazly, and M. Wink. 1997. Allelochemical activities of pyrrolizidine alkaloids: interactions with neuroreceptors and acetylcholine related enzymes. J Chem Ecol 23:399–416.

Sherratt T.N. 2003. State-dependent risk-taking by predators in systems with defended prey. Oikos 103:93–100.

Skelhorn J. and C. Rowe. 2007. Predators’ toxin burdens influence their strategic decisions to eat toxic prey. Curr Biol 17:1479–1483.

Skelhorn J., C.G. Halpin, and C. Rowe. 2016. Learning about aposematic prey. Behav Ecol 27:955–964.

Spinelli J.B. and M.C. Haigis. 2018. The multifaceted contributions of mitochondria to cellular metabolism. Nat Cell Biol 20:745–754.

Stier A., P. Bize, B.Y. Hsu, and S. Ruuskanen. 2019. Plastic but repeatable: rapid adjustments of mitochondrial function and density during reproduction in a wild bird species. Biol Lett 15:20190536.

Svensson L. 1992. Identification guide to European passerines. 4th ed. British Trust for Ornithology, Stockholm.

Timm K., K. Van Oers, and V. Tilgar. 2018. SERT gene polymorphisms are associated with risk-taking behaviour and breeding parameters in wild great tits. J Exp Biol 22: jeb171595.

Trigo J.R. 2011. Effects of pyrrolizidine alkaloids through different trophic levels. Phytochem Rev 10:83–98.

Von Nickisch-Rosenegk E. and M. Wink. 1993. Sequestration of pyrrolizidine alkaloids in several arctiid moths (Lepidoptera: Arctiidae). J Chem Ecol 19:1889–1903.

White I.N.H., A.R. Mattocks, and W.H. Butler. 1973. The conversion of the pyrrolizidine alkaloid retrorsine to pyrrolic derivatives in vivo and in vitro and its acute toxicity to various animal species. Chem Biol Interactions 6:207–218.

Wickham H. 2016. ggplot2: Elegant graphics for data analysis. 2nd ed. Springer international publishing, New York.

Wiedenfeld H. and J. Edgar. 2011. Toxicity of pyrrolizidine alkaloids to humans and ruminants. Phytochem Rev 10:137–151.

Winters A.E., J. Lommi, J. Kirvesoja, O. Nokelainen, and J. Mappes. 2021. Multimodal aposematic defenses through the predation sequence. Front Ecol Evol 9:657740.

Xu J., W. Wang, X. Yang, A. Xiong, L. Yang, and Z. Wang. 2019. Pyrrolizidine alkaloids: An update on their metabolism and hepatotoxicity mechanism. Liver Res 3:176–184.

Yang M., J. Ma, J. Ruan, Y. Ye, P.P.C. Fu, and G. Lin. 2019. Intestinal and hepatic biotransformation of pyrrolizidine alkaloid N-oxides to toxic pyrrolizidine alkaloids. Arch Toxicol 93:2197–2209.

