## Appendix for "The effect of the consumption of toxic prey on avian predator physiology"

### 1. *Arctia plantaginis* life-history traits

After the moth rearing was completed, we tested the effects of PA diet treatment on developmental time from larva to pupa (days), pupation time from pupa to eclosion (days) and pupal weight (mg). We used linear mixed-effects models to analyse larval developmental time and pupal weight, with treatment (PA/control), sex and pupation date (continuous) as fixed effects and family as a random effect to account for genetic variation. For pupation duration, family was not included because the random effect variance was effectively zero, so a linear model was adopted with treatment, sex and pupation date as fixed effects.

In total, 930 larvae pupated, but only 851 were used in the analysis (PA: n = 434, control: n = 417). These were the individuals that subsequently eclosed, allowing us to determine their sex. When analysing pupation time, we excluded four individuals with very short (< 5 days, n = 3) or long (18 days, n = 1) pupation times, as these were likely to be recording errors. In addition, one individual did not have an eclosion date recorded, and one was missing weight (final sample size for pupation time: n = 846, for weight: n = 850). For the analysis of developmental time from larvae to pupa, the response variable (time in days) was ln-transformed to improve normality. When analysing pupal weight and pupation time, there was heteroscedasticity between males and females, so we allowed different variances for each sex using the *nlme* package (Pinheiro and Bates 2025). There was also very minor heteroscedasticity between treatments in the developmental time model, but allowing different variances for each treatment did not improve model fit, so we used the simpler model.

Larvae fed on the PA diet (groundsel, *Senecio vulgaris*) had accelerated development (estimate = -0.073 ± 0.010, $t$ = -7.362, $p$ < 0.001) and pupated approximately four days faster than larvae fed on the control diet (dandelion only, Fig. A1a). There was also a tendency for males to develop faster than females, but this was not statistically significant (estimate = -0.017 ± 0.010, $t$ = -1.675, $p$ = 0.094). Both females (estimate = 18.194 ± 4.026, $t$ = 4.519, $p$ < 0.001) and males (estimate = 5.920 ± 2.728, $t$ = 2.170, $p$ = 0.030) had higher pupal weight when they were reared on the PA diet compared to the control diet, with a stronger effect in females (treatment × sex: estimate = -12.274 ± 4.856, $t$ = -2.527, $p$ = 0.012, Fig. A1b). Including family as a random effect improved model fit for both weight ($\sigma^2$ = 144.135) and larval development ($\sigma^2$ =0.009).

The time it took for pupae to emerge into adult *A.plantaginis* was not affected by diet treatment (estimate = -0.116 ± 0.074, $t$ = -1.566, $p$ = 0.12, Fig. A1c). However there was a significant sex effect (estimate = 0.871 ± 0.074, $t$ = 11.822, $p$ < 0.001), with males taking 0.87 days longer than females to pupate, and later pupation in the season was associated with longer pupation time (estimate = 0.028 ± 0.004, $t$ = 7.361, $p$ < 0.001).

Finally, we employed linear mixed-effect models for a subset of male only *A.plantaginis* individuals to explore a possible colour morph effect that could influence the results (white morph: n = 331, yellow morph: n = 75). It was found that the adult male phenotype did not affect larval development (estimate = -0.027 ± 0.023 $t$ = -1.149, $p$ = 0.25), pupation time (estimate = 0.132 ± 0.159, $t$ = 0.830, $p$ = 0.41) or male weight (estimate = 1.584 ± 4.141, $t$ = 0.383, $p$ =0.70).

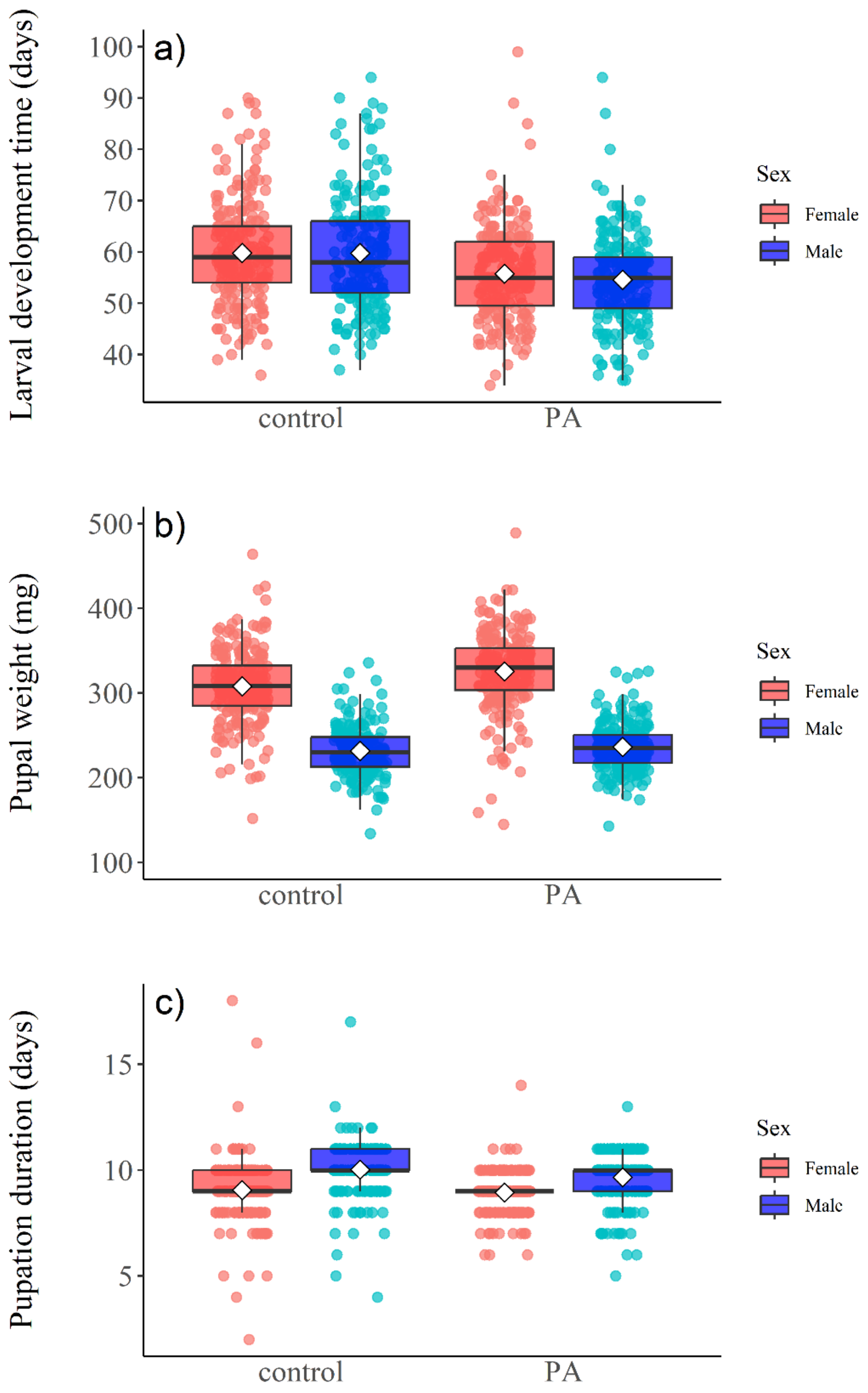

**Figure A1.** Differences between *A.plantaginis* reared on the control and PA diet in a) larval development time (days), b) pupal weight (mg), and c) pupal duration (days). Boxplots represent the median (horizontal line within the box), the first and third quartiles (lower and upper bounds of the box), and the whiskers indicate the values within 1.5 times the interquartile range. Circles represent individual values for each sample and white diamonds indicate the mean for each group.

### 2. Extraction and quantification of pyrrolizidine alkaloids

The method for the analysis of PAs in moths comprised a sample treatment procedure using acidic extraction and cleanup by solid phase extraction (SPE). Moths were first freeze-dried and homogenized, making a pool of each of the four treatment groups (see main text). The samples (20 mg of the homogenized powder, three technical replicates from each treatment pool) were then extracted with 0.05M sulfuric acid by sonicating them for 15 minutes in 40°C water, and then centrifuging for 10 minutes (4000 rpm). Strata SCX SPE cartridges were used for clean-up, and they were preconditioned with 2 mL methanol followed by 2 mL 0.05M sulphuric acid. After sample application, the cartridges were washed with 2 mL water, 2 mL methanol, vacuum dried for 2 min, and then 1 mL ethyl acetate was added. For elution of PAs, 2 mL of a solvent mixture containing ammonia and methanol – 6/94 v/v was added. The solvent was then evaporated under nitrogen at 40°C and reconstituted in methanol/water 50/50 v/v before LC-MS/MS analysis.

Reference standards with purities ranging from 95 to 99% were purchased from Phytolab Scientific (Tianjin, China). These standards included monocrotaline, retrorsine, senecionine, senecionine N-oxide, seneciphylline, and seneciphylline N-oxide. Monocrotaline was not detected in any of the samples, so it is not presented in the results.

The LC-MS/MS analysis was performed using Agilent 1290 ultra-high pressure LC system coupled to Agilent 6460 triple quadrupole mass spectrometer (Agilent Technologies, Waldbronn, Germany), operated in positive electrospray ionization with dynamic multiple reaction monitoring (MRM), acquiring specific product ions for each PA compound.

Chromatographic separation was performed using an Agilent InfinityLab Poroshell 120 EC-C18 column (100 mm × 2.1 mm (ID), particle size 2.7 μm), maintained at a constant temperature of 30°C. The sample injection volume was 3 μL. The mobile phase consisted of water containing 0.025% formic acid and 5 mM ammonium formate (solvent A), and methanol containing 0.025% formic acid and 5 mM ammonium formate (solvent B). Gradient elution was used at a flow rate of 0.3 mL/min. The gradient program was as follows: The initial eluent composition was 95% A, which was maintained for 3 minutes after injection; a linear gradient continued to 50% A over 12 minutes; held at 50% A for 2.5 minutes; the gradient continued to 28% A over 2.5 minutes; held for 0.5 minutes and then re-equilibrated to 95% A for 3.5 minutes. The analysis time was 21.5 minutes per sample.

Mass spectrometric detection was performed using an Agilent 6460 Triple Quadrupole equipped with an electrospray ionization (ESI) source utilizing Agilent Jet Stream Technology. The ESI source parameters were: sheath gas temperature at 400 °C, sheath gas flow at 10 L/min, drying gas temperature at 220 °C, gas flow at 10 L/min, nebulizer pressure at 30 psi, capillary voltage at 2500 V, and nozzle voltage at 1500 V. The mass spectrometer was operated in dynamic multiple reaction monitoring (dMRM) mode, monitoring two specific mass transitions per PA compound for accurate quantification and confirmation. The fragment voltage (FV), cell accelerator voltage (CAV), and collision energy (CE) for each PA compound are presented in Table S1. The CAV parameter was uniformly set at 7.

**Table A1**. Mass spectrometer parameters for each PA compound.

| Analyte | Rt (min) | Precursor ion $[M{+}H]^+$ (*m/z*) | Product ion Quantifier (*m/z*) | FV (V) | CE (eV) | Product ion Qualifier (*m/z*) | FV (V) | CE (eV) |
|---|---|---|---|---|---|---|---|---|
| Monocrotaline | 4.8 | 326 | 121 | 190 | 32 | 237 | 190 | 24 |
| Retrorsine | 8.8 | 352 | 120 | 190 | 32 | 138 | 190 | 32 |
| Senecionine | 10.3 | 336 | 120 | 190 | 28 | 138 | 190 | 32 |
| Senecionine *N*-oxide | 10.7 | 352 | 120 | 190 | 44 | 136 | 190 | 40 |
| Seneciphylline | 9.4 | 334 | 120 | 190 | 24 | 138 | 190 | 32 |
| Seneciphylline *N*-oxide | 9.7 | 350 | 120 | 190 | 36 | 136 | 190 | 40 |

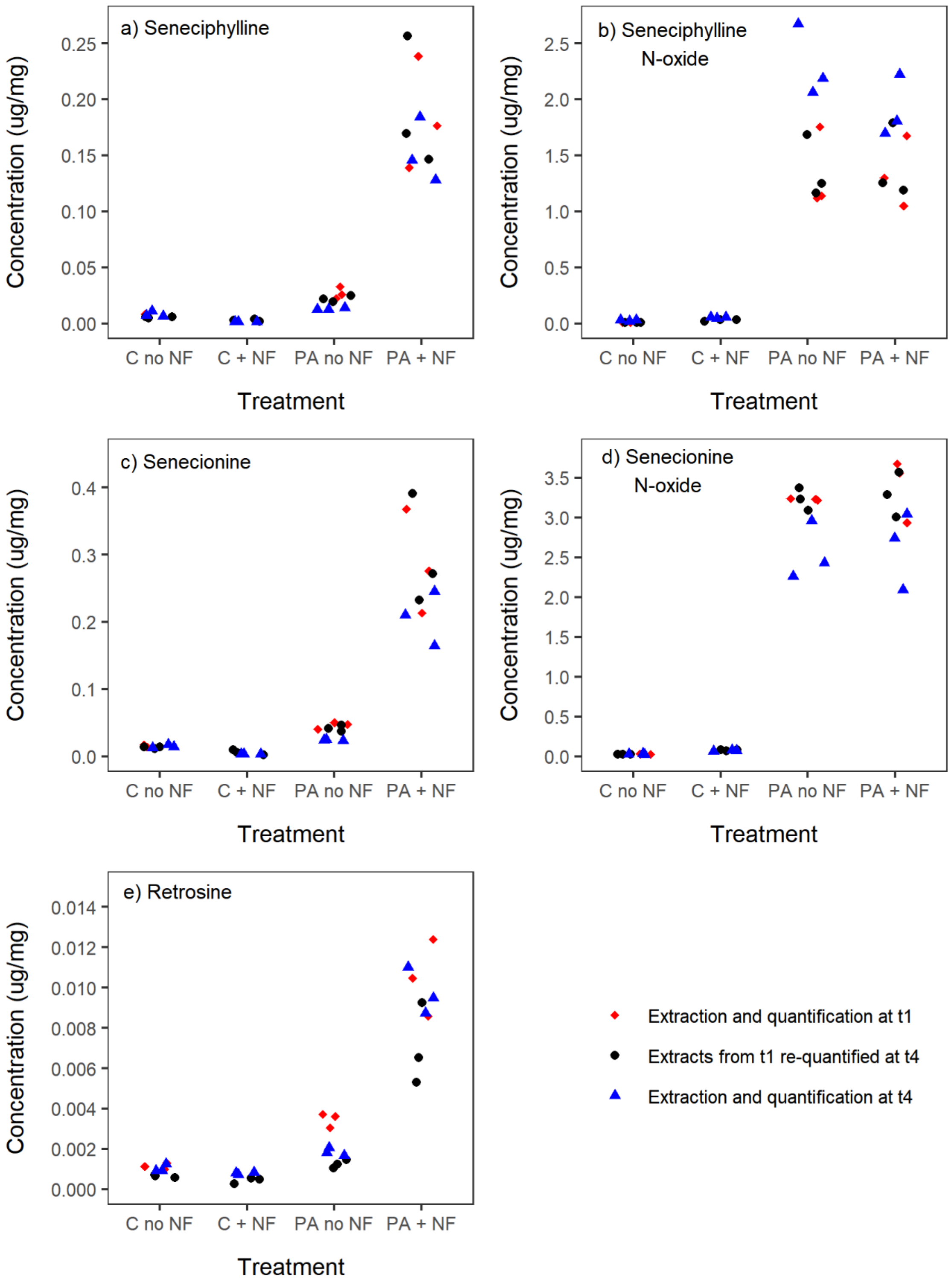

**Figure A2**. Quantified PA concentrations for a) seneciphylline, b) seneciphylline N-oxide, c) senecionine, d) senecionine N-oxide, and e) retrorsine. The graph shows the results from the extractions (three technical replicates from each treatment group) and quantifications performed at the beginning (t1, red diamonds) and at the end of the experiment (t4, blue triangles, and the results from the re-measurement of the t1 extracts at t4 (black circles). The quantifications of C + NF at t1 are excluded as they were outliers and likely to be a measurement error. Note the different y axis scale in each graph.

### 3. Full protocols for physiological assays

*Corticosterone concentration*

Frozen faecal samples from each treatment (n = 67) were first freeze-dried for 48 h using a Christ Alpha 1-4 LD Plus Lyophilizer. Then, 500 µL of ethanol was added to 50 mg of dried samples and shaken vigorously for 30 min using Qiagen TissueLyserII (25 Hz). The samples were then centrifuged for 15 min at 5000 rpm, and 420 µL of supernatant was dried under nitrogen for one hour. The dried extracted samples were stored at -20°C until analysis. Before the immunoassay, the sample was reconstituted in 12.5 µL of ethanol and 240 µL of assay buffer. 100 µL were then re-diluted into 140µL of the assay buffer.

Samples were analysed using the Arbor Assays DetectX Corticosterone EIA kit (Ann Arbor, MI, USA). To summarise, samples were added to wells coated with anti-goat antibody, and a corticosterone-peroxidase conjugate was added. After incubating for one hour, the plate was washed, and the substrate was added. This substrate reacts with the bound corticosterone-peroxidase conjugate. After a final short incubation (30 min), the reaction was stopped, and the plates were read using the Tecan Spark multimode microplate reader at 450nm. An additional standard 0 (10000 pg/mL) was added to the assay, due to the high corticosterone concentration of our samples (based on pilot tests). This ensured that the samples were in the linear part of the standard curve. All samples were analysed in duplicates. The extraction efficiency was estimated by spiking a sample with 1 µL of stock corticosteroid solution (concentration) and by performing 1:2, 1:4 and 1:8 dilutions. The average recovery of the serial dilutions was 108.45 %.

*Oxidative stress markers*

<u>Sample preparation</u>

Before analysing the biomarkers, we prepared and checked the protein concentration of the samples to assess their quality. All samples were handled on ice to avoid oxidation and degradation. First, 15 µL of each blood sample were diluted with 200 µL of 0.9% NaCl, to facilitate manipulation (Espin et al., 2017). Proteins were measured with Bradford method, using bovine serum albumin (BSA) as the standard. BSA standards (1 mg/mL) were pipetted in a 96-well plate, at volumes of 0, 2, 4, 6, 8 and 10 µL, serving as a standard range. A volume of 0.5 µL of each prepared blood sample was added per well. All samples, including standards, were measured in triplicate to calculate an intra-assay coefficient of variability (CV%, accepted values < 10 in all cases). Bio-Rad Protein Assay Dye Reagent Concentrate (Bio-Rad Laboratories, Cat #500–0006) diluted with MilliQ-water (1:5) was used as the reagent. After adding 200 µL of this mixture to each well, the plate was then incubated by shaking for 5 min at 600 rpm at room temperature. Absorbance was measured at 595 nm with the microplate reader (Tecan Spark multimode). Once the protein concentrations had been measured and validated, each sample was divided into three tubes with the quantities required for each analysis (12 µL for glutathione, 20 µL for GST and 100 µL for TBARS,). They were then immersed in liquid nitrogen before being stored at -80°C for future use.

GSH:GSSG

GSH concentration and GSH:GSSG ratio were analysed using the Arbor Assays DetectX Glutathione Fluorescent Detection Kit (Ann Arbor, MI, USA). Before starting, the samples were prepared by diluting 12 µL of blood from each bird (n = 66) with 48 µL of 5% SSA (Sulfosalicylic Acid + MQ-H2O), allowing the proteins to precipitate. The mixture was then vortexed and incubated for 10 min on ice, being vortexed three times during this period. The tubes were centrifuged for 10 min at 10000 g at 4 °C, and 30 µL of supernatant was collected and transferred to new tubes. Then, each sample was further diluted 1:5 in 120 µL of Assay Buffer (100 mM Na-phosphate buffer and 5 mM EDTA, pH 7.5) to give a final SSA concentration of 1%. Standards were prepared using Glutathione Standard (Sigma Chemicals, USA) diluted in Sample Diluent (100 mM Na-phosphate + 5 mM EDTA + 1% SSA). A volume of 50 µL of each standard was placed in the wells of a 96-well plate, as well as 50 µL of Sample Diluent for the blank. Then 50 µL of each prepared blood sample was added, and 25 µL of ThioStar was added to each well. The plate was then incubated at room temperature for 15 min. After this time, fluorescence intensity was measured at 510 nm with excitation at 390 nm using a microplate reader (Tecan Spark multimode). This measurement was used to estimate the amount of free GSH in the samples. Then, 25 µL of Reaction Mixture (Nicotinamide adenine dinucleotide phosphate (NADPH) concentrate (4 mM) + Glutathione reductase Concentrate (8 U ml-1 GR) + Assay Buffer) was added to each well, and the plate was left to incubate for another 15 min at room temperature. Fluorescence was measured at the same wavelengths, estimating the total amount of GSH (GSH + GSSG). All samples were measured in duplicates (accepted CV% < 15).

GST

To test the GST activity, 2 µL of prepared blood sample was pipetted in the wells of a 384-well plate. This was done in triplicates for each sample (n = 65). Just before taking the measurements, a solution of reaction mix (4.9 mL GST buffer (DPBS buffer = 1x Dulbecco's phosphate buffered Saline), 50 µL CDNB (1-Chloro-2,4-dinitrobenzene, 100 mM), 50 µL GSH (diluted in MQ-H2O, 200mM)) was prepared, and 48 µL of this solution was added to each well. The measurements were carried out using the microplate reader (Tecan Spark multimode) by measuring the absorbance of each plate 25 times at 340 nm. A control sample (made from a mixture of blood samples from randomly selected individuals) was used in each measurement to quantify inter-plate precision (CV% = 28.4).

Lipid peroxidation

TBARS analysis followed the procedure from Espin et al. (2017). Nine standards were prepared from MDA stock (Malonaldehyde bis (dimethyl acetal) 99%, 417 µM, diluted with ethanol), giving concentrations ranging from 0 to 2.085 nmol/µL. Then, a 1:100 solution of Butylated hydroxytoluene 2% (BHT diluted with ethanol) and TBARS reagent (15% Trichloroacetic acid, 0.25 N Hydrochloric acid, 0.375% 2-Thiobarbituric acid) was prepared and 200 µL were added to each tube. After vortexing, the samples were incubated for 30 min at 90°C on a heating block. The tubes were then placed on ice for 10 min to stop the reaction.

After this period, samples were centrifuged for 15 min at 6°C at 2100 g. Once centrifuged, 60 µL of supernatant from each sample was pipetted into a 96-well plate. Each sample was plated in triplicate to obtain an intra-assay coefficient of variability (accepted CV% < 15). The fluorescence intensity was then measured at an excitation/emission of 540/590 nm with a microplate reader (Tecan Spark multimode). Two of the three plates included a control (made from a mixture of blood samples) to quantify inter-plate precision (CV% = 21). This control was only used on the last two plates out of three due to a handling error.

*Serotonin concentration*

Before the 5-HT analysis, plasma was diluted in a 1:20 ratio, using 5µL of blood and 95uL of assay buffer. Samples from each treatment (n = 42) were analysed using a commercially available Serotonin ELISA kit (ADI-900-175, Enzo Ltd., Farmingdale, NY), following the manufacturer's instructions. To sum up briefly, standards and samples were added to wells coated with a goat anti-rabbit immunoglobulin G antibody. A solution of serotonin conjugated to alkaline phosphatase was then added, followed by a solution of rabbit polyclonal antibody to serotonin. After a two hour incubation, the plate was washed, leaving only bound serotonin. P-nitrophenyl phosphate substrate was then added. This generates a yellow colour when catalysed by the serotonin conjugate. After a final incubation (one hour), a stop solution was added. Plates were then read at 405nm using the Tecan Spark multimode microplate reader. Samples were analysed in duplicates when possible (i.e. if there was enough plasma).

### 4. The number of birds used in each analysis

**Table A2:** Final sample size in each statistical model after excluding outliers and samples with high CV%. The table shows the number of adults ( > 1 year old) and juveniles (< 1 year old), and males and females in each bird diet treatment.

| | C +no NF | | | | C + NF | | | | PA + no NF | | | | PA + NF | | | | Total |
|---|---|---|---|---|---|---|---|---|---|---|---|---|---|---|---|---|---|
| | ♂ AD | ♂ JUV | ♀ AD | ♀ JUV | ♂ AD | ♂ JUV | ♀ AD | ♀ JUV | ♂ AD | ♂ JUV | ♀ AD | ♀ JUV | ♂ AD | ♂ JUV | ♀ AD | ♀ JUV | |
| Total | N = 20 | | | | N = 20 | | | | N = 20 | | | | N = 20 | | | | 80 |
| | 6 | 5 | 3 | 6 | 3 | 7 | 4 | 6 | 7 | 5 | 0 | 8 | 5 | 6 | 1 | 8 | |
| Consumption latency | N = 16 | | | | N = 16 | | | | N = 17 | | | | N = 16 | | | | 65 |
| | 4 | 4 | 3 | 5 | 3 | 6 | 3 | 4 | 6 | 3 | 0 | 8 | 5 | 5 | 1 | 5 | |
| Weight change | N = 11 | | | | N = 10 | | | | N = 11 | | | | N = 8 | | | | 40 |
| | 3 | 1 | 3 | 4 | 1 | 4 | 2 | 3 | 4 | 1 | 0 | 6 | 3 | 2 | 1 | 2 | |
| Corticosterone | N = 16 | | | | N = 16 | | | | N = 13 | | | | N = 14 | | | | 59 |
| | 5 | 5 | 2 | 4 | 3 | 6 | 3 | 4 | 3 | 4 | 0 | 6 | 4 | 4 | 1 | 5 | |
| GST activity | N = 11 | | | | N = 8 | | | | N = 10 | | | | N = 12 | | | | 41 |
| | 2 | 3 | 1 | 5 | 1 | 1 | 3 | 3 | 3 | 2 | 0 | 5 | 4 | 2 | 0 | 6 | |
| GSH | N = 17 | | | | N = 15 | | | | N = 16 | | | | N = 15 | | | | 63 |
| | 6 | 4 | 2 | 5 | 2 | 5 | 2 | 6 | 5 | 4 | 0 | 7 | 5 | 3 | 1 | 6 | |
| GSH:GSSG | N = 16 | | | | N = 15 | | | | N = 16 | | | | N = 13 | | | | 60 |
| | 5 | 4 | 2 | 5 | 2 | 5 | 2 | 6 | 5 | 4 | 0 | 7 | 5 | 2 | 1 | 5 | |
| TBARS | N = 14 | | | | N = 15 | | | | N = 16 | | | | N = 14 | | | | 59 |
| | 5 | 4 | 2 | 3 | 2 | 5 | 3 | 5 | 6 | 3 | 0 | 7 | 5 | 3 | 1 | 5 | |
| Mitochondrial metabolism | N = 8 | | | | N = 12 | | | | N = 13 | | | | N = 10 | | | | 43 |
| | 4 | 1 | 2 | 1 | 1 | 5 | 2 | 4 | 5 | 1 | 0 | 7 | 2 | 5 | 1 | 2 | |
| Serotonin | N = 10 | | | | N = 10 | | | | N = 7 | | | | N = 10 | | | | 37 |
| | 4 | 2 | 3 | 1 | 2 | 5 | 1 | 2 | 2 | 2 | 0 | 3 | 3 | 2 | 1 | 4 | |

### 5. Full model outputs and model selections

**Table A3.** Linear mixed effects model explaining the time to consume the moth pellet (min), with a) full output of the final model, and b) model comparison. One bird that was missing body condition measure was excluded when comparing models (n = 64), but it was included in the final model (n = 65). Bird identity was included in all models as a random effect. BC = body condition, MD = manipulation day.

a) final model summary

| | **Effect** | **SE** | **t** | **P** |
|---|---|---|---|---|
| Intercept | 188.694 | 13.370 | 14.113 | < 0.001 |
| Treatment: C + NF | 15.038 | 15.497 | 0.970 | 0.34 |
| Treatment: PA + no NF | 47.088 | 15.304 | 3.077 | 0.003 |
| Treatment: PA + NF | 54.987 | 15.518 | 3.543 | < 0.001 |
| Manipulation day | -11.484 | 2.195 | -5.232 | < 0.001 |

b) model comparison

| **Alternative models** | **df change** | $\chi^2$ | **P** | **AIC** |
|---|---|---|---|---|
| ~ treatment + MP (final model) | | | | 4306.7 |
| ~ treatment + MP + age | 1 | 0.610 | 0.43 | 4308.1 |
| ~ treatment + MP + age + BC | 1 | 0.012 | 0.91 | 4310.1 |
| ~ treatment + MP + age + BC + sex | 1 | 0.032 | 0.86 | 4312.1 |
| ~ treatment × MP + age + BC + sex | 3 | 5.750 | 0.12 | 4312.3 |

**Table A4.** Linear model explaining birds' weight change (g) after the diet manipulation (difference between capture weight and weight after the six day manipulation), with a) full output of the final model and b) model comparison (n = 40).

a) final model summary

| | **Effect** | **SE** | **t** | **P** |
|---|---|---|---|---|
| Intercept | -1.629 | 0.249 | -6.550 | < 0.001 |
| Treatment: C + NF | 0.048 | 0.360 | 0.133 | 0.90 |
| Treatment: PA + no NF | 0.291 | 0.352 | 0.827 | 0.41 |
| Treatment: PA + NF | 0.002 | 0.383 | 0.004 | 0.997 |

b) model comparison

| **Alternative models** | **df** | **F** | **P** | **AIC** |
|---|---|---|---|---|
| ~ treatment (final model) | 36 | | | 103.90 |
| ~ treatment + sex | 35 | 0.789 | 0.38 | 104.98 |
| ~ treatment + sex + age | 34 | 0.152 | 0.70 | 106.81 |

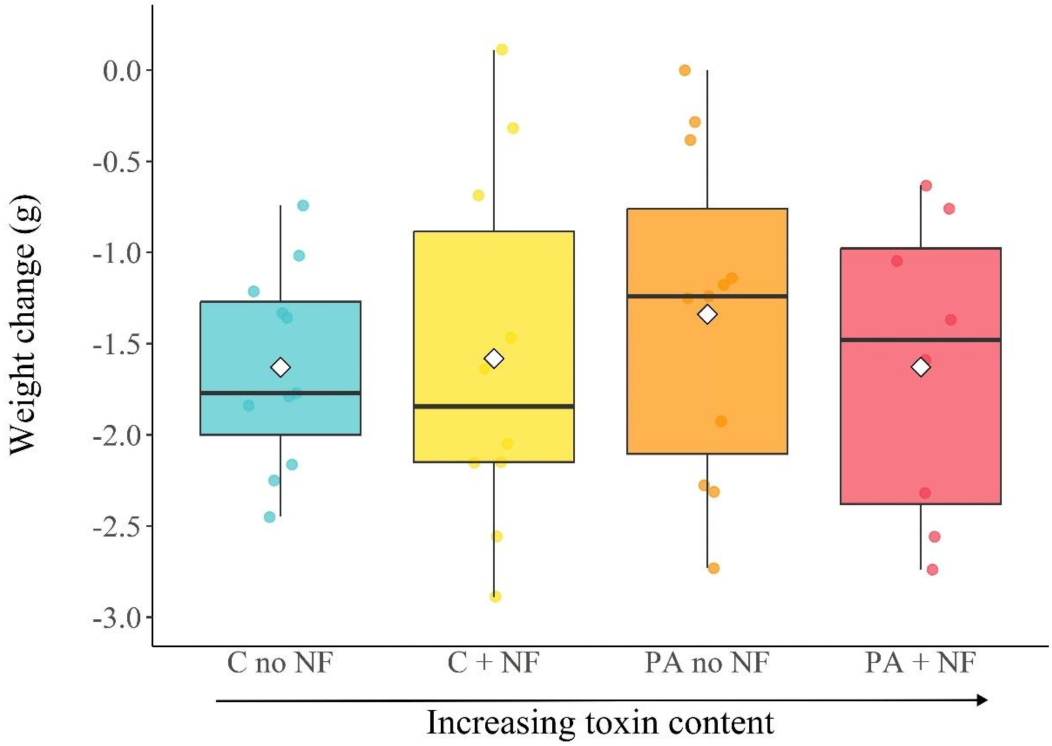

**Figure A3.** Weight change (g) after the six-day diet manipulation in each bird diet treatment. Boxplots represent the median (horizontal line within the box), the first and third quartiles (lower and upper bounds of the box), and the whiskers indicate the values within 1.5 times the interquartile range. Circles represent individual values for each sample and white diamonds indicate the mean for each treatment.

**Table A5.** Linear model explaining corticosterone concentration (ng/g, $\log_{10}$ transformed), with a) full output of the final model and b) model comparison. Two birds that were missing body condition measure were excluded when comparing models (n = 57), but they were included in the final model (n = 59). BC = body condition, PN = assay plate number.

a) final model summary

| | **Effect** | **SE** | **t** | **P** |
|---|---|---|---|---|
| Intercept | 2.832 | 0.058 | 48.542 | < 0.001 |
| Treatment: C + NF | -0.033 | 0.066 | -0.497 | 0.62 |
| Treatment: PA + no NF | -0.059 | 0.070 | -0.846 | 0.40 |
| Treatment: PA + NF | -0.014 | 0.069 | -0.208 | 0.84 |
| PN: second plate | -0.138 | 0.051 | -2.730 | 0.009 |

b) model comparison

| **Alternative models** | **df** | **F** | **P** | **AIC** |
|---|---|---|---|---|
| ~ treatment + PN (final model) | 52 | | | -22.33 |
| ~ treatment + PN + BC | 51 | 0.159 | 0.69 | -20.52 |
| ~ treatment + PN + BC + sex | 50 | 0.096 | 0.76 | -18.63 |
| ~ treatment + PN + BC + sex + age | 49 | 0.002 | 0.96 | -16.63 |

**Table A6.** Linear model explaining MDA concentration (nmol/mg protein) in the TBARS assay, with a) full output of the final model and b) model comparison. Two birds that were missing body condition measures were excluded when comparing models (n = 57), but they were included in the final model (n = 59). BC = body condition, PN = assay plate number.

a) final model summary

| | Effect | SE | t | P |
|---|---|---|---|---|
| Intercept | 0.0067 | 0.001 | 7.163 | < 0.001 |
| Treatment: C + NF | -0.0018 | 0.001 | -1.736 | 0.09 |
| Treatment: PA + no NF | 0.0007 | 0.001 | 0.655 | 0.51 |
| Treatment: PA + NF | -0.0002 | 0.001 | -0.221 | 0.83 |
| PN: second plate | 0.0039 | 0.001 | 4.223 | < 0.001 |
| PN: third plate | 0.0077 | 0.001 | 8.273 | < 0.001 |

b) model comparison

| Alternative models | df | F | P | AIC |
|---|---|---|---|---|
| ~ treatment + PN (final model) | 51 | | | -496.27 |
| ~ treatment + PN + age | 50 | 0.098 | 0.30 | -495.55 |
| ~ treatment + PN + age + BC | 49 | 0.253 | 0.61 | -493.85 |
| ~ treatment + age + BC + sex | 48 | 0.142 | 0.71 | -492.02 |

**Table A7.** Linear model explaining GST activity (pmol/min/mg of protein, ln-transformed), with a) full output of the final model, and b) model comparison. One bird that was missing body condition measure was excluded when comparing models (n = 40), but it was included in the final model (n = 41). BC = body condition, PN = assay plate number.

a) final model summary

| | Effect | SE | t | P |
|---|---|---|---|---|
| Intercept | -10.275 | 0.150 | -68.686 | < 0.001 |
| Treatment: C + NF | 0.068 | 0.231 | 0.297 | 0.77 |
| Treatment: PA + no NF | 0.035 | 0.217 | 0.160 | 0.87 |
| Treatment: PA + NF | 0.294 | 0.207 | 1.419 | 0.16 |

b) model comparison

| Alternative models | df | F | P | AIC |
|---|---|---|---|---|
| ~ treatment (final model) | 36 | | | 61.75 |
| ~ treatment + PN | 33 | 1.505 | 0.23 | 62.48 |
| ~ treatment + PN + age | 32 | 0.662 | 0.42 | 63.65 |
| ~ treatment + PN + age + sex | 31 | 0.938 | 0.34 | 64.44 |
| ~ treatment + PN + age + sex + BC | 30 | 0.487 | 0.49 | 65.79 |

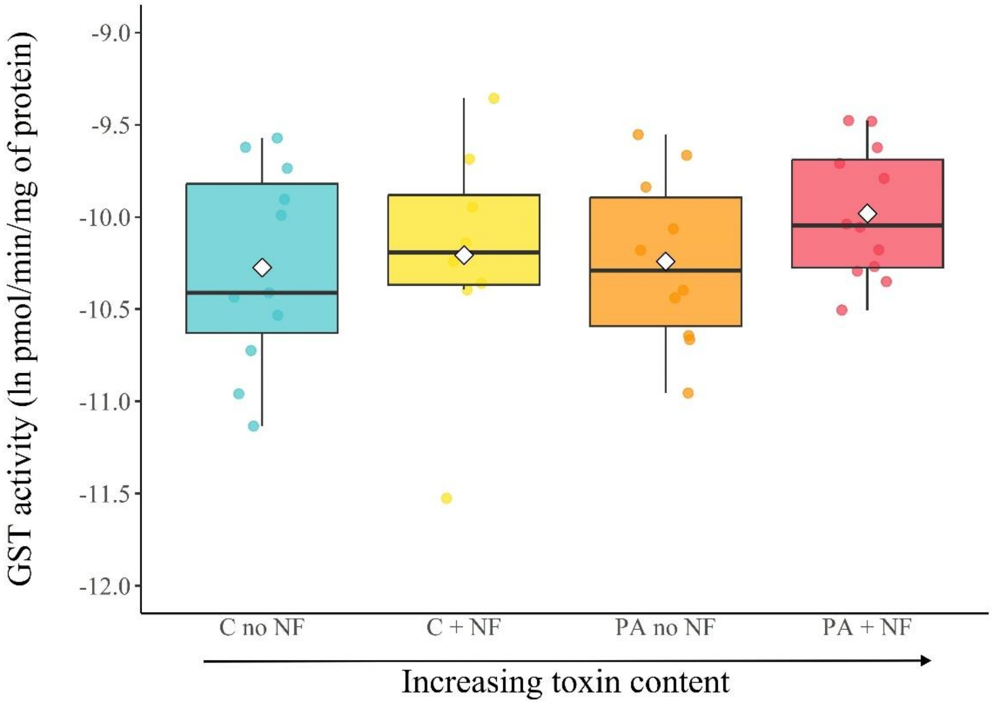

**Figure A4.** GST activity (pmol/min/mg of protein, ln-transformed) in each bird diet treatment. Boxplots represent the median (horizontal line within the box), the first and third quartiles (lower and upper bounds of the box), and the whiskers indicate the values within 1.5 times the interquartile range. Circles represent individual values for each sample and white diamonds indicate the mean for each treatment.

**Table A8.** Linear model explaining GSH concentration (nmol/mg protein, ln-transformed), with a) full output of the final model, and b) model comparison. Two birds that were missing body condition measures were excluded when comparing models (n = 61), but they were included in the final model (n = 63). BC = body condition, PN = assay plate number.

a) final model summary

| | **Effect** | **SE** | **t** | **P** |
|---|---|---|---|---|
| Intercept | -1.753 | 0.081 | -21.654 | < 0.001 |
| Treatment: C + NF | -0.032 | 0.118 | -0.267 | 0.79 |
| Treatment: PA + no NF | 0.038 | 0.116 | 0.323 | 0.75 |
| Treatment: PA + NF | -0.007 | 0.118 | -0.057 | 0.96 |

b) model comparison

| **Alternative models** | **df** | **F** | **P** | **AIC** |
|---|---|---|---|---|
| ~ treatment (final model) | 57 | | | 45.88 |
| ~ treatment + PN | 56 | 0.162 | 0.69 | 47.70 |
| ~ treatment + PN + BC | 55 | 2.053 | 0.16 | 47.39 |
| ~ treatment + PN + BC + age | 54 | 0.221 | 0.64 | 49.14 |
| ~ treatment + PN + BC + age + sex | 53 | 0.117 | 0.73 | 51.01 |

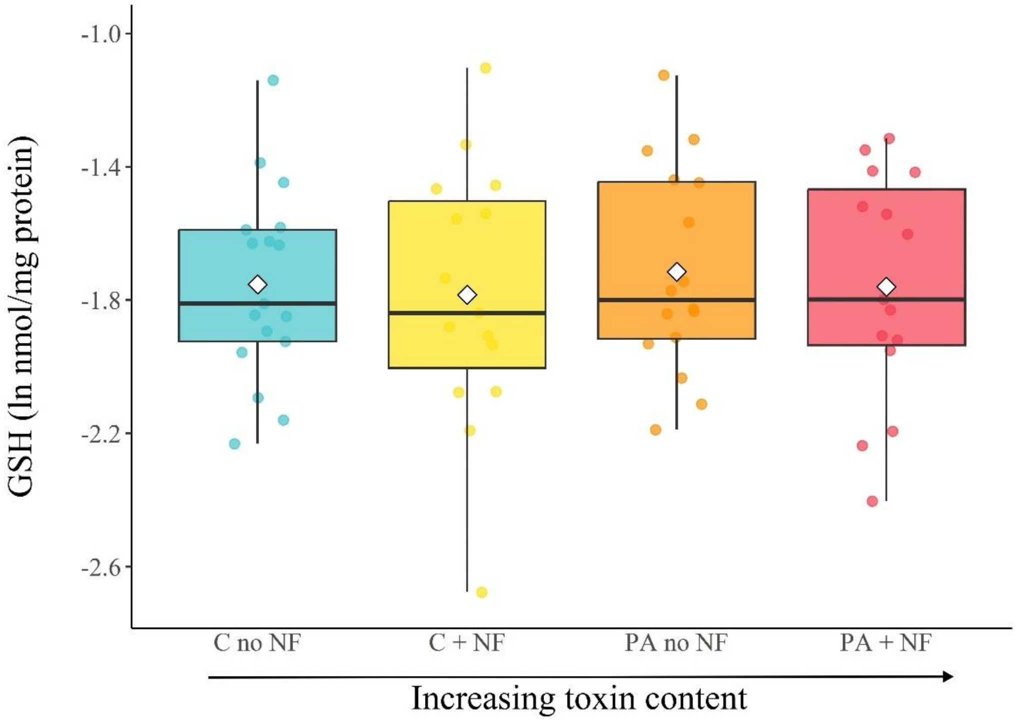

**Figure A5.** GSH concentration (nmol/mg of protein, ln-transformed) in each bird diet treatment. Boxplots represent the median (horizontal line within the box), the first and third quartiles (lower and upper bounds of the box), and the whiskers indicate the values within 1.5 times the interquartile range. Circles represent individual values for each sample and white diamonds indicate the mean for each treatment.

**Table A9.** Linear model explaining the GSH:GSSG ratio (ln-transformed), with a) full output of the final model and b) model comparison. Two birds that were missing body condition measures were excluded when comparing models (n = 58), but they were included in the final model (n = 60). BC = body condition, PN = assay plate number.

a) final model summary

| | **Effect** | **SE** | **t** | **P** |
|---|---|---|---|---|
| Intercept | 0.899 | 0.132 | 6.839 | < 0.001 |
| Treatment: C + NF | -0.047 | 0.175 | -0.265 | 0.79 |
| Treatment: PA + no NF | -0.227 | 0.175 | -1.293 | 0.20 |
| Treatment: PA + NF | 0.027 | 0.182 | 0.148 | 0.88 |
| PN: second plate | 0.488 | 0.132 | 3.691 | < 0.001 |

b) model comparison

| **Alternative models** | **df** | **F** | **P** | **AIC** |
|---|---|---|---|---|
| ~ treatment + PN (final model) | 53 | | | 89.13 |
| ~ treatment + PN + BC | 52 | 1.235 | 0.27 | 89.75 |
| ~ treatment + PN + BC + age | 51 | 1.128 | 0.29 | 90.46 |
| ~ treatment + PN + BC + age + sex | 50 | 0.135 | 0.72 | 92.30 |

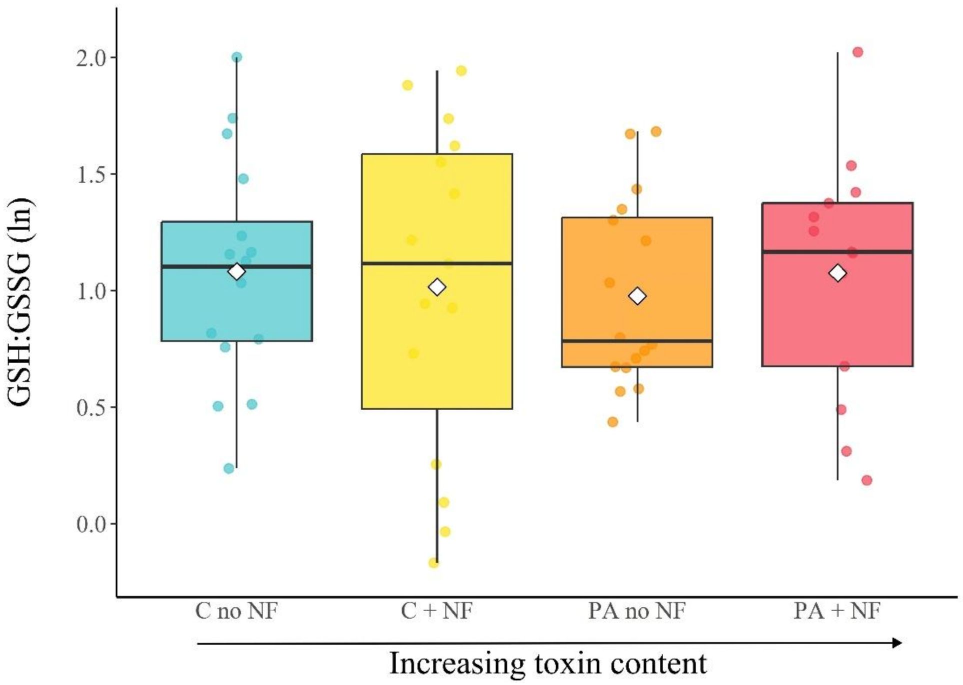

**Figure A6.** GSH:GSSG ratio (ln-transformed) in each bird diet treatment. Boxplots represent the median (horizontal line within the box), the first and third quartiles (lower and upper bounds of the box), and the whiskers indicate the values within 1.5 times the interquartile range. Circles represent individual values for each sample and white diamonds indicate the mean for each treatment.

**Table A10.** Linear model explaining routine mitochondrial respiration (pmol/s/mL), with a) full output of the final model and b) model comparison. Two birds that were missing body condition measures were excluded when comparing models (n = 41) but they were included in the final model (n = 43). BC = body condition, CC = cell count.

a) final model summary

| | **Effect** | **SE** | **t** | **P** |
|---|---|---|---|---|
| Intercept | 0.020 | 0.071 | 0.277 | 0.78 |
| Treatment: C + NF | 0.090 | 0.043 | 2.075 | 0.045 |
| Treatment: PA + no NF | 0.108 | 0.043 | 2.516 | 0.016 |
| Treatment: PA + NF | 0.063 | 0.046 | 1.364 | 0.18 |
| Sex: male | 0.048 | 0.030 | 1.632 | 0.11 |
| Cell count | 0.031 | 0.011 | 2.742 | 0.009 |

b) model comparison

| **Alternative models** | **df** | **F** | **P** | **AIC** |
|---|---|---|---|---|
| ~ treatment + CC | 36 | | | -69.38 |
| ~ treatment + sex + CC (final model) | 35 | 2.977 | 0.09 | -70.73 |
| ~ treatment + sex + CC + BC | 34 | 1.248 | 0.27 | -70.21 |
| ~ treatment + sex + CC + BC + sex + age | 33 | 0.799 | 0.38 | -69.20 |

**Table A11.** Linear model explaining oxidative phosphorylation (OXPHOS), with a) full output of the final model and b) model comparison. Two birds that were missing body condition measures were excluded when comparing models (n = 41) but they were included in the final model (n = 43). BC = body condition, CC = cell count.

a) final model summary

| | Effect | SE | t | P |
|---|---|---|---|---|
| Intercept | 0.041 | 0.004 | 9.208 | < 0.001 |
| Treatment: C + NF | 0.006 | 0.006 | 0.998 | 0.33 |
| Treatment: PA + no NF | 0.002 | 0.006 | 0.296 | 0.77 |
| Treatment: PA + NF | 0.004 | 0.006 | 0.706 | 0.48 |
| Age: juvenile | 0.007 | 0.004 | 1.593 | 0.12 |

b) model comparison

| Alternative models | df | F | P | AIC |
|---|---|---|---|---|
| ~ treatment | 37 | | | -235.09 |
| ~ treatment + age (final model) | 36 | 2.338 | 0.14 | -235.81 |
| ~ treatment + age + BC | 35 | 0.990 | 0.33 | -235.02 |
| ~ treatment + age + BC + sex | 34 | 0.089 | 0.77 | -233.13 |
| ~ treatment + age + BC + sex + CC | 33 | 0.001 | 0.97 | -231.13 |

**Table A12.** Linear model explaining FCR ROUTINE/(CI+II) ratio, with a) full output of the final model and b) model comparison. Two birds that were missing body condition measures were excluded when comparing models (n = 41) but they were included in the final model (n = 43). BC = body condition, CC = cell count.

a) final model summary

| | Effect | SE | t | P |
|---|---|---|---|---|
| Intercept | 0.089 | 0.031 | 2.896 | 0.006 |
| Treatment: C + NF | 0.034 | 0.019 | 1.838 | 0.074 |
| Treatment: PA + no NF | 0.069 | 0.018 | 3.729 | < 0.001 |
| Treatment: PA + NF | 0.026 | 0.020 | 1.303 | 0.20 |
| Sex: male | 0.042 | 0.013 | 3.271 | 0.002 |
| Cell count | 0.008 | 0.005 | 1.681 | 0.10 |

b) model comparison

| Alternative models | df | F | P | AIC |
|---|---|---|---|---|
| ~ treatment + sex | 36 | | | -139.53 |
| ~ treatment + sex + CC (final model) | 35 | 3.161 | 0.08 | -141.26 |
| ~ treatment + sex + CC + BC | 34 | 0.213 | 0.65 | -139.52 |
| ~ treatment + sex + CC + BC + age | 33 | 0.043 | 0.84 | -137.57 |

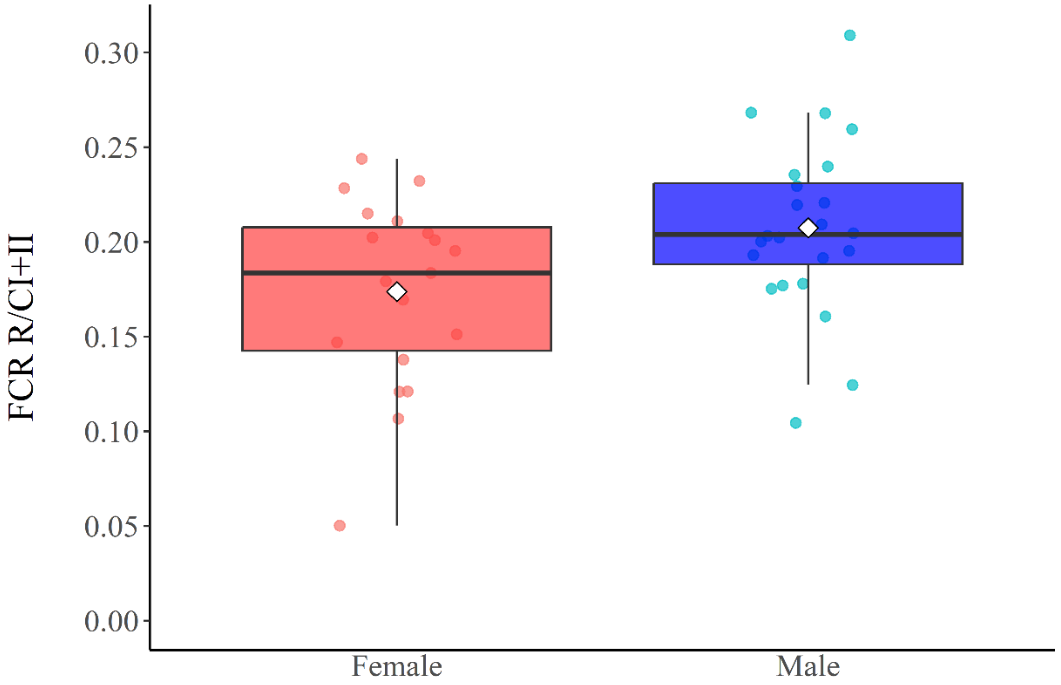

**Figure A7:** FCR ROUTINE/(CI+II) ratio in females and males. Boxplots represent the median (horizontal line within the box), the first and third quartiles (lower and upper bounds of the box), and the whiskers indicate the values within 1.5 times the interquartile range. Circles represent individual values for each sample and white diamonds indicate the mean for each treatment.

**Table A13.** Linear model explaining serotonin concentration (ng/mL, $log_{10}$ transformed) with a) full output of the final model and b) model comparison (n = 37). BC = body condition.

a) final model summary

| | **Effect** | **SE** | **t** | **P** |
|---|---|---|---|---|
| Intercept | 0.396 | 1.216 | 0.326 | 0.75 |
| Treatment: C + NF | -0.068 | 0.207 | -0.327 | 0.75 |
| Treatment: PA + no NF | 0.026 | 0.240 | 0.106 | 0.92 |
| Treatment: PA + NF | -0.052 | 0.207 | -0.254 | 0.80 |
| BC | 0.118 | 0.066 | 1.801 | 0.08 |

b) model comparison

| **Alternative models** | **df** | **F** | **P** | **AIC** |
|---|---|---|---|---|
| ~ treatment | 33 | | | 52.67 |
| ~ treatment + BC (final model) | 32 | 3.179 | 0.08 | 51.10 |
| ~ treatment + BC + sex | 31 | 1.059 | 0.31 | 51.83 |
| ~ treatment + BC + sex +age | 30 | 0.312 | 0.58 | 53.4 |

### Appendix references

Espín, S., P. Sánchez-Virosta, and A.J. García-Fernández. 2017. A microplate adaptation of the thiobarbituric acid reactive substances assay to determine lipid peroxidation fluorometrically in small sample volumes. Rev Toxicol 34:94–98.

Pinheiro, J., D. Bates, and R Core Team. 2025. nlme: Linear and nonlinear mixed effects models. R package version 3.1-168.
